# RNF43 inhibits WNT5A driven signaling and suppresses melanoma invasion

**DOI:** 10.1101/2021.02.08.430210

**Authors:** Tomasz Radaszkiewicz, Michaela Nosková, Kristína Gömöryová, Olga Vondálová Blanářová, Katarzyna Anna Radaszkiewicz, Markéta Picková, Ráchel Víchová, Tomáš Gybel’, Karol Kaiser, Lucia Demková, Lucia Kučerová, David Potěšil, Zbyněk Zdráhal, Karel Souček, Vítězslav Bryja

## Abstract

RNF43 is a E3 ubiquitin ligase and known negative regulator of WNT/β-catenin signaling. We demonstrate that RNF43 is also regulator of noncanonical WNT5A-induced signaling in human cells. Analysis of the RNF43 interactome using BioID and immunoprecipitation showed that RNF43 can interact with the core receptor complex components dedicated to the noncanonical Wnt pathway such as ROR1, ROR2, VANGL1 and VANGL2. RNF43 triggers VANGL2 ubiquitination and proteasomal degradation and clathrin-dependent internalization of ROR1 receptor. This activity of RNF43 is physiologically relevant and blocks pro-metastatic WNT5A signaling in melanoma. RNF43 inhibits responses to WNT5A, which results in the suppression of invasive properties of melanoma cells. Furthermore, RNF43 prevented WNT5A-assisted development of resistance to BRAF V600E inhibitor. In line with these findings, *RNF43* expression decreases during melanoma progression and RNF43-low patients have worse prognosis. We conclude that RNF43 is a newly discovered negative regulator of WNT5A-mediated biological responses that desensitizes cells to WNT5A.

## Introduction

Ubiquitination is a post-translational modification (PTM) based on the addition of evolutionary conserved protein ubiquitin (Ub) to the lysine residue(s) of the modified protein (Hershko and Ciechanover, 1998). Ubiquitination controls turnover, activation state, cellular localization, and interactions of target proteins. Undoubtfully, it is a process, which has a direct impact on various aspects of cell biology (Rape, 2018). Ubiquitination requires sequential activation of ubiquitin, its transfer to the carrier protein and subsequent linkage reaction with the substrate lysine residues. This last step, mediated by the E3 ubiquitin-protein ligases (E3s), determines target specificity.

Ring Finger protein 43 (RNF43) is a E3 ubiquitin ligase with single transmembrane domain from the PA-TM-RING family. RNF43 and its close homolog Zinc and Ring Finger 3 (ZNRF3), act as negative regulators of the Wnt/β-catenin signaling pathway (Koo *et al*, 2012; Hao *et al*, 2012). Wnt/β-catenin signaling is an evolutionary conserved pathway and a crucial regulator of embryonal development and tissue homeostasis. RNF43 and ZNRF3 control via regulation of Wnt/β-catenin multiple processes including liver zonation (Planas-Paz et al., 2016), limb specification (Szenker-Ravi et al., 2018) and mammalian sex determination (Harris et al., 2018). Mechanistically, RNF43 and ZNRF3 ubiquitinate plasma membrane Wnt receptors called Frizzleds (FZDs) and a co-receptor Low-density Lipoprotein Receptor-related Protein 6 (LRP6), which results in their internalization and degradation (Hao et al., 2012; Koo et al., 2012). Therefore, cells become less sensitive or insensitive to Wnt ligands. Activity of RNF43/ZNRF43 is regulated by secreted proteins from R-spondin (RSPO) family (Kazanskaya et al., 2004; Kim et al., 2008, 2006, 2005; Nam et al., 2007, 2006; Peng et al., 2013; Xie et al., 2013) that trigger internationalization of RNF43/ZNRF3 and function as physiologically relevant activators of Wnt/β-catenin pathway (Binnerts et al., 2007; Carmon et al., 2011; de Lau et al., 2011; Hao et al., 2016, 2012; Jiang et al., 2015; Koo et al., 2012; Zebisch et al., 2013; Zebisch and Jones, 2015).

Because deregulation of Wnt/β-catenin pathway promotes tumor formation (Lim and Nusse, 2013; van Kappel and Maurice, 2017; Wiese et al., 2018), RNF43/ZNRF3 can act as tumor suppressors. Indeed, mutation or inactivation of *RNF43*/*ZNRF3* lead to the oncogenic activation of Wnt signaling and associates with colorectal, liver, gastric, endometrial, ovarian and pancreatic cancers (Bond et al., 2016; Eto et al., 2018; Giannakis et al., 2014; Jiang et al., 2013; Jo et al., 2015; Niu et al., 2015; Planas-Paz et al., 2016; Ryland et al., 2013; Spit et al., 2020; Tsukiyama et al., 2020).

Some members of the Wnt family – such as WNT5A and WNT11 - preferentially activate downstream signaling that is distinct from Wnt/β-catenin pathway and is referred to as β-catenin-independent or noncanonical Wnt pathway (Pandur *et al*, 2002; Humphries & Mlodzik, 2018; VanderVorst *et al*, 2019; Andre *et al*, 2015). Noncanonical Wnt pathway shares some features with the Wnt/β-catenin pathway – such as requirement for FZD receptors, Dishevelled (DVL) phosphoprotein and Casein Kinase 1 (CK1) – but clearly differs in others. In mammalian noncanonical pathway Receptor Tyrosine Kinase Like Orphan Receptor 1 (ROR1) and ROR2 act as primary (co-)receptors (in contrast to LRP5/6 that have this role in the Wnt/β-catenin pathway) and four-transmembrane Vang-like protein 1 (VANGL1) and VANGL2 participate on the signal transduction (Asem et al., 2016; VanderVorst et al., 2019). This signaling axis is also referred to as Planar Cell Polarity Pathway (PCP) and its activation leads to the changes in the actin cytoskeleton dynamics, facilitating i.e. polarized cell migration (Andre et al., 2015; Janovská and Bryja, 2017; Kaucká et al., 2015; Weeraratna et al., 2002).

FZD receptors, the best-defined targets of RNF43/ZNRF3, are shared among all Wnt pathways and their endocytosis and/or degradation have the potential, at least in theory, to prevent signaling by any Wnt ligands. So far, however, there are is no systematic study addressing the role of RNF43/ZNRF3 in the noncanonical Wnt signaling in mammals. On the other side there are several hints that suggest that such possibility is feasible. Secreted inhibitor of RNF43/ZNRF3 called r-spondin 3 (RSPO3), potentiated noncanonical PCP pathway in *Xenopus* in a Wnt5a and Dishevelled-dependent manner (Glinka et al., 2011; Ohkawara et al., 2011). In mouse embryos *Znrf3* knockout caused open neural tube defects, which is a common consequence of the Wnt/PCP signaling disruption (Hao et al., 2012). Other report showed similar phenotype in *Xenopus* embryos after *Rnf43* mRNA injection (Tsukiyama et al., 2015). And finally, in *Caenorhabditis elegans*, the homolog of RNF43 and ZNRF3 called plr-1 was shown to control not only surface localization of frizzled, but also proteins related to mammalian noncanonical Wnt co-receptors ROR1/2 and RYK (Moffat et al., 2014). However, it is worth to underline that RSPO family homologs are absent in *C. elegans* (Lebensohn and Rohatgi, 2018), so the mode of action of RNF43/ZNRF3 in worm might be different than in mammalian cells.

In this study, we have directly addressed the role of RNF43 in the WNT5A-induced signaling. We demonstrate that RNF43 controls noncanonical Wnt pathway similarly to Wnt/β-catenin pathway. We demonstrate that RNF43 is a relevant inhibitor of pro-metastatic WNT5A signaling in melanoma where it prevents both WNT5A-induced invasive behavior and WNT5A-assisted development of resistance to B-RAF inhibitors.

## Results

### RNF43 inhibits WNT5A driven noncanonical Wnt signaling pathway

In order to test whether or not RNF43/ZNRF3 controls noncanonical Wnt signaling, we have decided to study T-REx 293 cells. T-REx 293 cells secrete endogenous WNT5A that constitutively activates noncanonical Wnt pathway – this can be demonstrated by the CRISPR/Cas9-mediated knockout of *WNT5A* (Kaiser et al., 2020). Removal of the endogenous WNT5A in T-REx 293 cells is sufficient to eliminate activation of readouts of WNT5A signaling such as phosphorylation of ROR1, DVL2 and DVL3 that can be monitored as the decrease in the phosphorylation-mediated electrophoretic mobility shifts (Fig. 1A). Such autocrine WNT5A signaling is promoted by the inhibition of endogenous RNF43/ZNRF3 by RSPO1 treatment (Fig. 1B, compare lane 1 and 2) and inhibited by RNF43 overexpression under the tetracycline (Tet) controlled promoter (TetON) or by block of WNT secretion using porcupine inhibitor Wnt-C59 (Fig. 1B). To confirm that the effects are indeed caused by block of WNT5A signaling, T-REx 293 cells pre-treated with Wnt-C59 and as such unable to produce Wnt ligands, were stimulated with the increasing doses of recombinant WNT5A. As shown in Fig. 1C, overexpression of RNF43 completely blocked signaling induced by recombinant WNT5A. Altogether, this demonstrates that RNF43 has the potential to block WNT5A signaling in mammalian cells.

**Figure 1.**
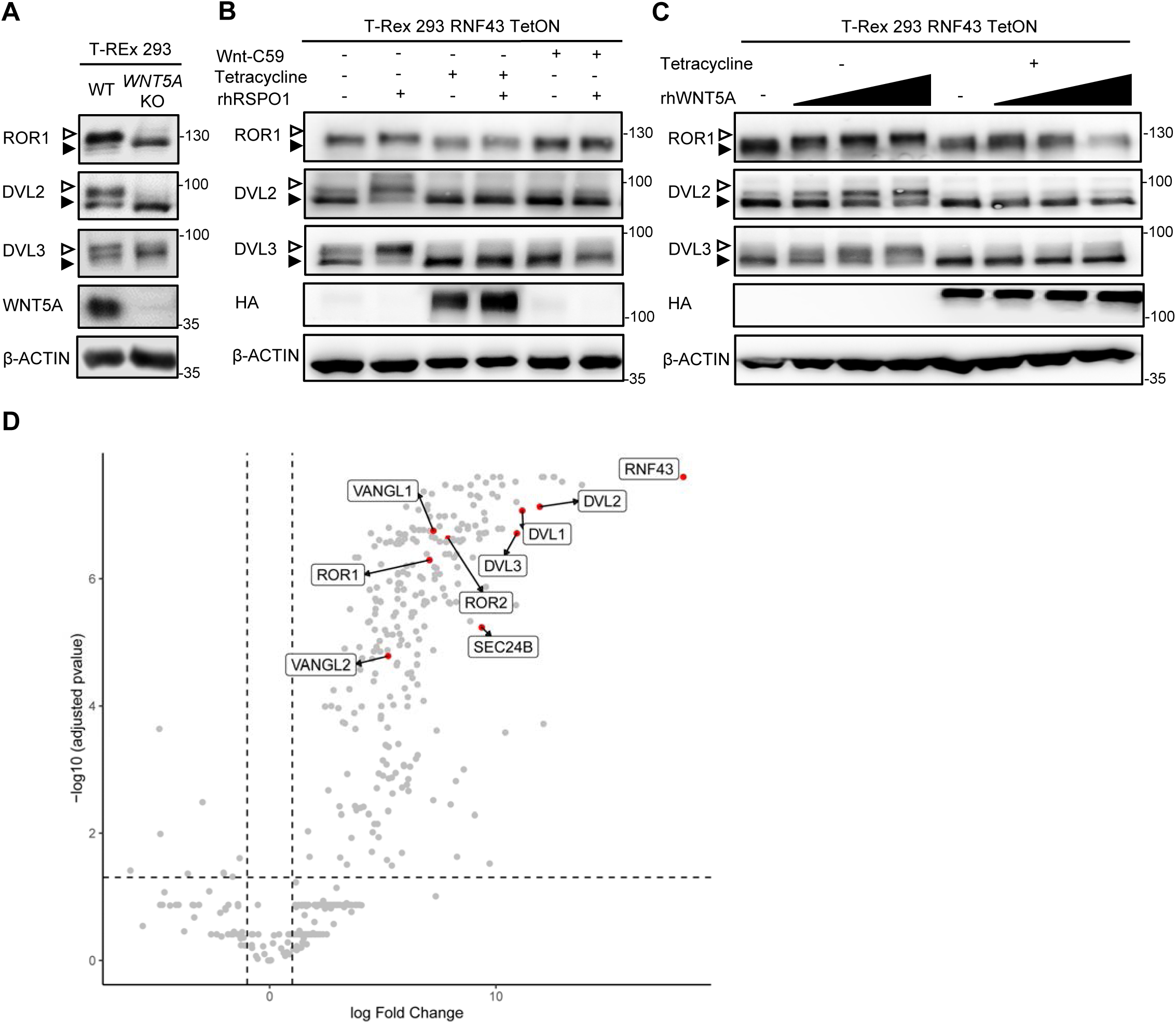
RNF43 interactome is enriched with the Wnt Planar Cell Polarity pathway components. **A**. Western blot analysis of T-REx 293 WNT5A KO and parental cells. Phosphorylation dependent shifts of endogenous ROR1, DVL2 and DVL3 were suppressed upon WNT5A loss. Signal of β-actin serves as a loading control **B**. Western blot showing activation of the noncanonical Wnt pathway components: ROR1, DVL2 and DVL3 (arrowheads) upon rhRSPO1 overnight treatment. Tetracycline forced RNF43 overexpression (as visualized by HA tag specific antibody) suppressed this effect. Inhibition of Wnt ligands secretion by the porcupine inhibitor Wnt-C59 shows dependency of the rhRSPO1 mediated effect on endogenous Wnt ligands; representative blots from N=3. **C**. Western blot analysis of cellular responses to the increasing doses of rhWNT5A. ROR1 shift and phosphorylation of DVL2 and DVL3 (arrowheads) were inhibited upon tetracycline induced RNF43-HA-BirA* overexpression. All samples were treated with Wnt-C59 to ascertain assay specificity to the exogenous rhWNT5A, N=3. **D**. Volcano plot of the RNF43 interactome identified by BioID and subsequent mass spectrometric detection (see M&M for details). Significantly enriched proteins annotated as the components of the noncanonical Wnt signaling pathway are highlighted. Full list of BioID-based identified interactors of RNF43 is present in the Figure 1 Supplementary table 1 and GO terms enrichment analysis in the Figure 1 Supplementary table 2.

### RNF43 physically interacts with key proteins from noncanonical WNT pathway

To address the molecular mechanism of RNF43 action in the noncanonical Wnt pathway we decided to describe RNF43 interactome by the proximity-dependent biotin identification (BioID) (Roux et al., 2012), which was already successfully applied in the challenging identification of E3s substrates (Coyaud et al., 2015; Deshar et al., 2016). We have exploited our recently published dataset (Spit et al., 2020) based on T-REx 293 TetON cells that inducibly expressed RNF43 fused C-terminally (intracellularly) with BirA* biotin ligase. Several core proteins of the noncanonical Wnt signaling pathway – namely ROR1, ROR2, VANGL1, VANGL2, SEC24B and all three isoforms of DVL – were strongly and specifically biotinylated by RNF43-BirA* (Fig. 1D and Figure 1 Supplementary table 1). Furthermore, noncanonical Wnt pathway was significantly enriched also in the gene ontology (GO) terms (Figure 1 Supplementary table 2). Altogether, it suggests that RNF43 can at least transiently interact with multiple proteins involved in the Wnt/Planar Cell Polarity pathway, including essential receptor complex components from the ROR, DVL and VANGL protein families.

To validate the protein-protein interactions identified by BioID, we performed a series of co-immunoprecipitation (co-IP) and co-localization experiments (Fig. 2 and Figure 2 figure supplement 1). We have focused on the interactions of RNF43 with ROR1/ROR2 and with VANGL1/VANGL2 mainly because these interactions are novel and at the same time highly relevant for the noncanonical Wnt pathway. RNF43 co-immunoprecipitated with both VANGL2 (Fig. 2A) and VANGL1 (Figure 2 figure supplement 1A). More detailed analysis of VANGL2 showed co-localization of VANGL2 and RNF43 in the cell membrane (Fig. 2B, B’). RNF43 also efficiently pulled down ROR1 (Fig. 2C) and ROR2 (Figure 2 figure supplement 1B). Deletion of the cysteine rich domain (CRD) (ROR2, Figure 2 figure supplement 1B) had no impact on the amount of co-immunoprecipitated RNF43, which suggests that RNF43 primarily interacts with RORs intracellularly. Both ROR1/ROR2 co-localized with RNF43 at the level of plasma membrane (Fig. 2D, D’ and Figure 2 figure supplement 1C, C’). It was described that RORs and VANGLs also bind DVL (Gao et al., 2011; Mentink et al., 2018; Seo et al., 2017; Witte et al., 2010; Yang et al., 2017) and at the same time DVL proteins mediate ubiquitination of FZD receptors by RNF43 in the Wnt/β-catenin pathway (Jiang et al., 2015). To address whether DVL also acts as a physical link between RNF43 and the analyzed PCP proteins we performed the co-IP experiments with VANGL2 and ROR1 in the T-REx 293 cells lacking all free DVL isoforms (DVL triple knockout cells) (Paclíková et al., 2017). As shown in Figure 2 figure supplement 1D and E, RNF43 was able to bind both VANGL2 and ROR1 as efficiently as in the wild type cells (compare with Fig. 2A, D). In summary, our results indicate that RNF43 interacts, in a DVL-independent way, with PCP proteins from VANGL and ROR families.

**Figure 2.**
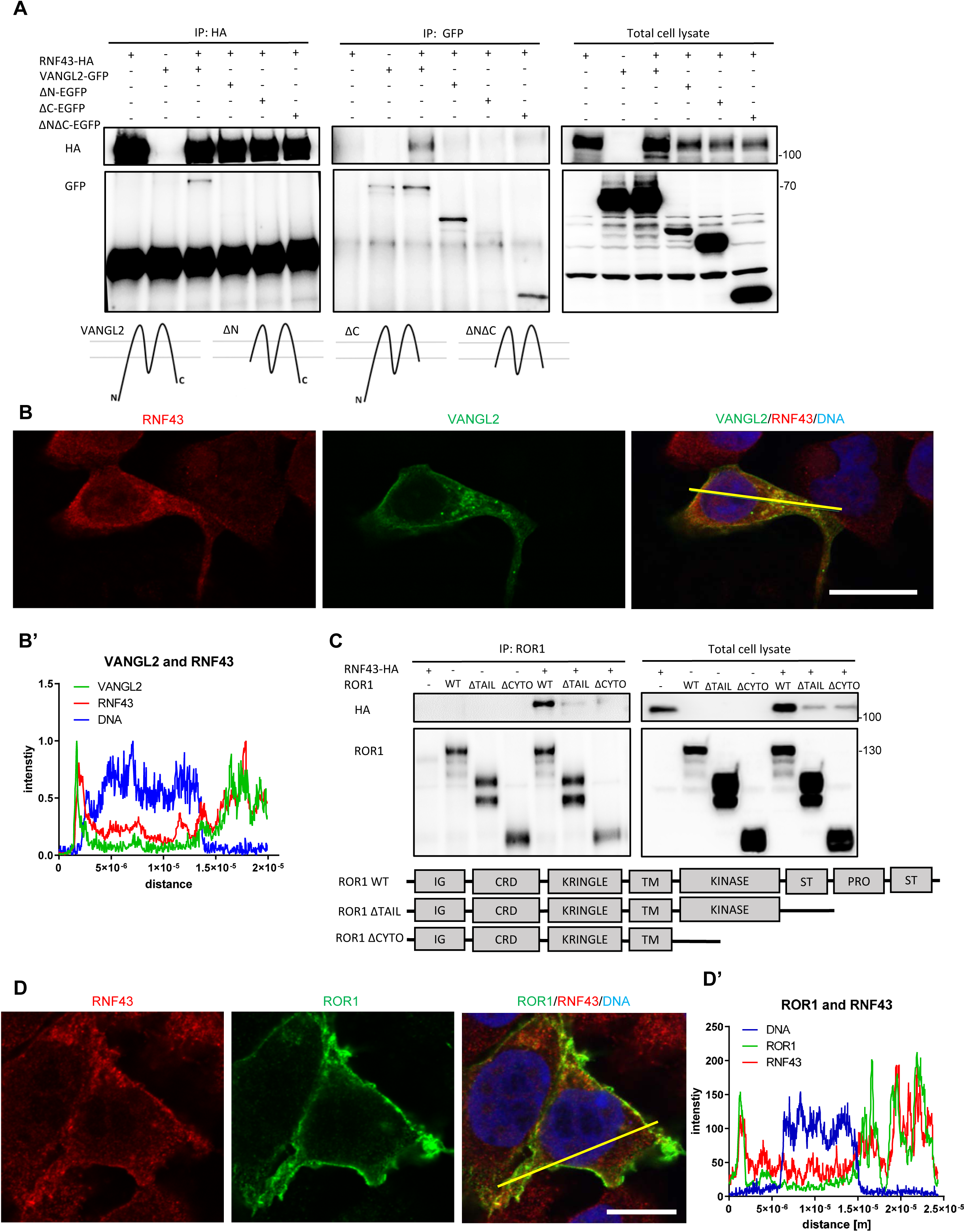
RNF43 interacts with Wnt/PCP components. **A**. RNF43 interacts with VANGL2, but not with its mutants lacking N- or C-termini. VANGL2-EGFP and its variants (schematized) were overexpressed with RNF43-HA in Hek293 T-REx cells, immunoprecipitated by anti-HA and anti-GFP antibodies and analyzed by Western blotting. Representative experiment from N=3. Scheme illustrates secondary structure of the wild type VANGL2 protein and its shortened variants used in this study. **B**., **B’**. RNF43 (anti-HA, red) colocalized with transiently expressed VANGL2 (GFP, green). Co-localization was analyzed utilizing histograms of red, green and blue channels signals along selection (yellow line) (B’). TO-PRO-3 Iodide was used to stain nuclei (blue). Scale bar: 25 μm. **C**. RNF43 binds to the ROR1 and deletion of the intracellular part of ROR1 disrupts this interaction. RNF43-HA was detected in the ROR1 pull down prepared from lysates of Hek293 T-REx cells overexpressing RNF43-HA and ROR1-V5, N=3. ROR1 wild type and truncated mutants are represented in the scheme. **D**., **D’**. RNF43 (anti-HA, red) colocalized with transiently expressed ROR1-V5 (anti-V5, green). Signals along selection (yellow line) were analyzed (D’). TO-PRO-3 was employed nuclei staining (blue). Scale bar:25 μm. RNF43 interactions with VANGL1 and ROR2 are studied in the Figure 2 figure supplement 1. Figure 2 Source Data contains raw data used in the B’ and D’.

### RNF43 ubiquitinates VANGL2 and triggers its degradation

Since RNF43 is an E3 ubiquitin ligase we next tested whether it can ubiquitinate its binding partners from the noncanonical Wnt pathway. Enzymatically inactive RNF43 Mut1 variant (Koo et al., 2012), served here as a negative control. Using His-ubiquitin pulldown assay, we were able to show that VANGL2 (Fig. 3A), as well as DVL1 and DVL2 (Figure 3 figure supplement 1A) were ubiquitinated when co-expressed with RNF43 but not with RNF43 Mut1. However, we were unable to detect RNF43-induced ubiquitination of ROR1 or ROR2 (negative data, not shown).

**Figure 3.**
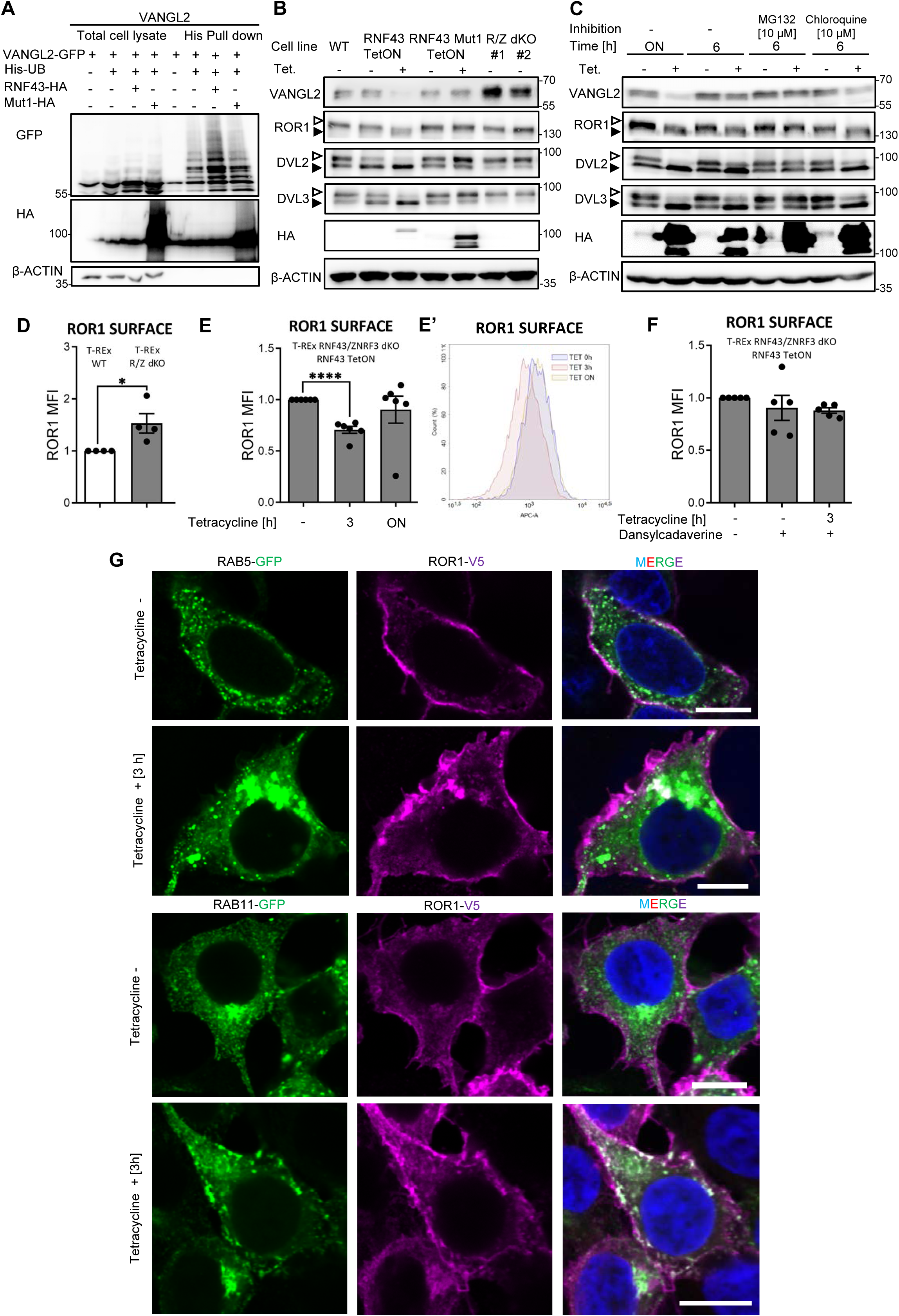
Mechanism of Wnt/PCP inhibition by RNF43. **A**. Hek293 T-REx cells were transfected with plasmid encoding His-tagged ubiquitin, VANGL2-GFP and HA-tagged wild type or Mut1 RNF43 constructs. Ubiquitinated proteins were enriched by by His pull down and analyzed by Western blotting. VANGL2 is ubiquitinylated by the E3 ubiquitin ligase RNF43, but not by its enzymatically inactive variant (RNF43Mut1). Representative experiment from N=3. RNF43-mediated ubiquitination of DVL1 and DVL2 together in the Figure 3 figure supplement 1. **B**. Tetracycline-induced overexpression of the wt RNF43 (HA), but not enzymatically inactive RNF43Mut1 (HA), decreased VANGL2 protein level and suppressed phosphorylation of ROR1 and DVL3 (open arrowheads). CRISPR/Cas9 derived RNF43/ZNRF3 (R/Z) dKO cell lines #1 and #2 displayed phenotype reversed to the RNF43 overexpression. Quantified in RNF43-mediated ubiquitination of DVL1 and DVL2 together in the Figure 3 figure supplement 1B, N=3. **C**. Inhibition of the proteasomal degradation pathway by MG132 (but not by lysosomal inhibitor chloroquine) blocked the RNF43 effects on ROR1, DVL2, DVL3 and VANGL2 as shown by the Western blotting analysis, N=3. **D**. Flow cytometric analysis of surface ROR1 in wild type (WT) and RNF43/ZNRF3 (R/Z) dKO cells; unpaired two-tailed t-test: p= 0.0298, N=4. ROR1 was stained using ROR1-APC conjugate on the not permeabilizated cells. Validation of the a-ROR1-APC antibody is shown in the Figure 3 figure supplement 1D. **E**., **E’**. Surface ROR1 levels upon 3h and overnight (ON) induction of RNF43 in RNF43 TetON RNF43/ZNRF3 dKO cells; unpaired t-test p< 0.0001, N=6. Representative histogram of ROR1-APC signal in the analyzed conditions is shown (E’). **F**. Dansylcadaverine, inhibitor of clathrin-mediated endocytosis, blocked the effect of RNF43 overexpression on surface ROR1, performed as in E; N=5. **G**. Immunofluorescence imaging showed enhanced ROR1(V5) colocalization with the marker of early endosomes RAB5 (GFP) after 3h tetracycline treatment in RNF43 TetON RNF43/ZNRF3 dKO cells. (bottom) RAB11 positive (GFP) recycling endosomes were recruited to the ROR1 (V5) at the plasma membrane after overnight tetracycline treatment. Cells were transfected, treated, fixed and stained. DNA was visualized by Hoechst 33342. Similar results were obtained for T-REx RNF43 TetON cell line (Figure 3 figure supplement 1E-G). Raw data used in the D, E and F are encolsed in the Figure 3 Source Data. Data presented in the B, E and F is presented in the Figure 3 figure supplement Source Data file.

Further analysis showed that overexpression of RNF43, but not its E3 ligase dead variant, decreased VANGL2 protein level (Fig. 3B, quantified in Figure 3 figure supplement 1B). Decrease in VANGL2 caused by RNF43 was accompanied by impeded phosphorylation of ROR1 (Fig. 3B) and DVL3 (Fig. 3B, Figure 3 figure supplement 1B). On the other side, two independent clones of cells deficient in both RNF43 and ZNRF3 (*RNF43*/*ZNRF3* dKO; *R*/*Z* dKO) showed higher VANGL2 levels and higher DVL phosphorylation (Fig. 3B, Figure 3 figure supplement 1B). Interestingly, treatment with proteasome inhibitor MG132 but not with autophagosome-lysosome inhibitor Chloroquine blocked these effects of RNF43 (Fig. 3C). This suggests that RNF43 action in noncanonical Wnt pathway depends on the proteasomal degradation pathway, which differs from the Wnt/β-catenin pathway, where RNF43 triggers FZD degradation via lysosomal pathway (Koo et al., 2012).

### RNF43 induces ROR1 endocytosis by a clathrin dependent pathway

ROR1 and ROR2 are the key receptors for WNT5A that we found to interact with RNF43 (Figs. 1 and 2). We thus speculated that RNF43 can regulate ROR1/ROR2 surface levels. T-Rex cells express dominantly ROR1 and indeed flow cytometric analysis demonstrated that cell lacking endogenous RNF43 and ZNRF3 have more ROR1 receptor on the surface than parental T-REx cells (Fig. 3D). The staining is specific as demonstrated by the validation of the ROR1-APC antibody in *ROR1* KO T-REx 293 cells (Figure 3 figure supplement 1C and D). When we introduced inducible RNF43 into *RNF43*/*ZNRF3* dKO T-REx cell line, we were able to rescue this phenotype and after three hours of tetracycline treatment we detected decreased surface ROR1 (Fig. 3E, E’). The overnight exposition to tetracycline had no significant effect (Fig. 3E, E’). Similar trends were observed for wild type T-REx 293 RNF43 TetON cells (Figure 3 figure supplement 1E, E’).

In our analysis of RNF43 interactors (Fig. 1D), we identified also multiple proteins involved in the endosomal transport. It included proteins involved in the clathrin endocytic pathway - STAM1, HRS, ZFYVE16, PICALM, NUMB, RAB11-FIP2 and subunits of the associated adaptor protein complexes AP-3 and AP-4 (Figure 1 Supplementary table 1) (Bache et al., 2003; Cullis et al., 2002; Hirst et al., 2013; Raiborg et al., 2001; Santolini et al., 2000; Seet and Hong, 2005; Tebar et al., 1999). Based on the BioID results analysis, we thus speculated that RNF43 may promote clathrin-mediated endocytosis of ROR1. Thus, we applied dansylcadaverine to block this pathway (Blitzer and Nusse, 2006). In agreement with our hypothesis, treatment with this inhibitor prevented RNF43-mediated effect on the ROR1 surface expression in both cell lines (Fig. 3F, Figure 3 figure supplement 1F).

To get a better insight into the mechanism of RNF43-induced internalization of ROR1, we analyzed the colocalization of ROR1 and RAB5 (marker of early endosomes) and RAB11 (marker of recycling endosomes) in T-REx 293 R/Z dKO RNF43 TetON (Fig. 3G and Figure 3 figure supplement 2) and T-REx 293 RNF43 TetON cells (Figure 3 figure supplement 1G and Figure 3 figure supplement 2). Hyperactivation of Rab5 by overexpression of wild-type Rab5 leads to the formation giant early endosomes (Bucci et al., 1992) where we observed ROR1/RAB5 co-localization after three hours of tetracycline treatment. The co-localization decreased after overnight exposition to tetracycline. RAB11^+^ endosomes were recruited to the ROR1 as well after RNF43 induction and RAB11 co-localized strongly with ROR1 even after ON treatment. We conclude that surface ROR1 is controlled by RNF43 via interference with RAB5 and RAB11 mediated endocytosis and vesicle recycling.

### RNF43 expression is decreased in human melanoma

Our data shown in Figs. 1-3 demonstrate that RNF43 can inhibit WNT5A-induced noncanonical signaling via downregulation of the receptor complexes. But is RNF43 capable to block WNT5A-induced biological processes? WNT5A signaling plays crucial role in melanoma, one of the most malignant tumor types. High expression of *WNT5A* in this cancer is a negative overall survival and positive metastasis formation factor (Da Forno et al., 2008; Luo et al., 2020; Weeraratna et al., 2002). Signaling cascade activated by WNT5A in melanoma drives epithelial– mesenchymal transition (EMT), resulting in the increased metastatic properties of melanoma cells *in vitro* and *in vivo* (Dissanayake et al., 2008, 2007; Sadeghi et al., 2018). In melanoma WNT5A acts through FZD and ROR1/ROR2 (O’Connell et al., 2010; Tiwary and Xu, 2016; Weeraratna et al., 2002). Importance of WNT5A driven signaling in melanoma is thus well recognized and melanoma represents probably the most characterized (and most clinically relevant) pathophysiological condition where noncanonical WNT5A signaling drives cell invasion and disease progression (Arozarena and Wellbrock, 2017a; Da Forno et al., 2008; Dissanayake et al., 2007; Lai et al., 2012; Liu et al., 2018; O’Connell et al., 2010, 2008; Weeraratna et al., 2002).

Interestingly, the *in silico* analysis of gene expression in melanoma (Talantov et al., 2005; Xu et al., 2008) showed that *RNF43* expression dramatically decreases between benign melanocytic skin nevus and cutaneous melanoma (Fig. 4A) (Talantov et al., 2005) and further between primary site and metastasis (Xu et al., 2008) (Fig. 4B). Importantly, analysis of other datasets (Anaya, 2016) showed that *RNF43* low melanoma patients have shorter overall survival (OS) (Fig. 4C). *ZNRF3* expression had no prognostic value (Figure 4 figure supplement 1D). Interestingly, expression of two genes encoding direct targets ubiquitinated by RNF43, namely *DVL3* and *VANGL1*, increased during melanoma progression (Figure 4 figure supplement 1A, C) and high expression in both cases correlates with bad prognosis and shorter overall survival (Fig. 4D, Figure 4 figure supplement 1 B). All these findings are in line with the hypothesis that RNF43 acts in melanoma as a tumor suppressor that restricts WNT5A-induced biological processes and gets silenced during melanoma progression.

**Figure 4.**
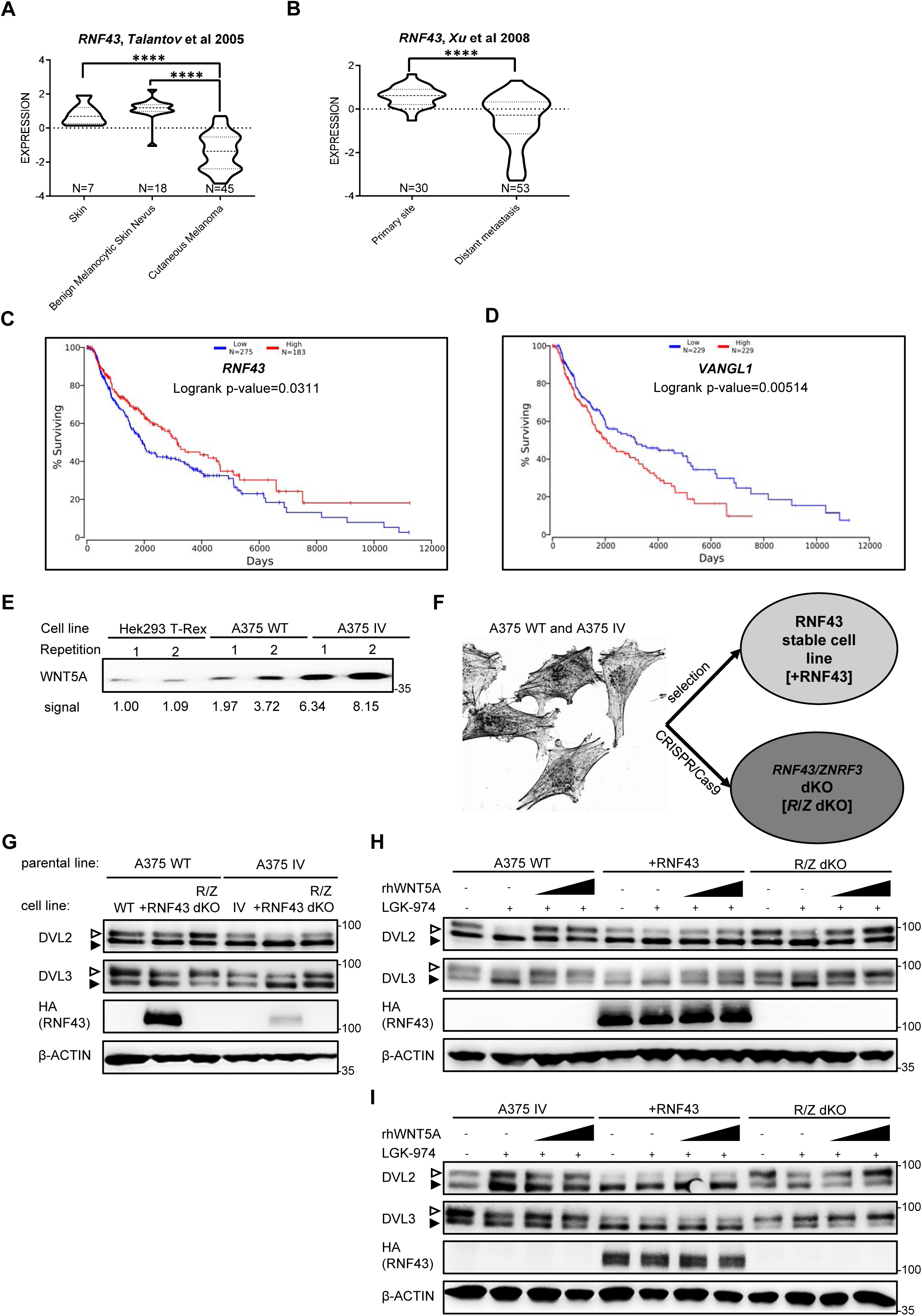
RNF43 in melanoma. **A**., **B**. *RNF43* expression is lower in melanoma when compared with the skin and benign melanocytic skin nevus (A) and in the case of distant metastasis compared to the primary tumors (B), unpaired two-tailed t-test: **** p<0.0001. **C**. *RNF43* expression is negative prognostic factor in melanoma. RNF43 low patients have shorter overall survival (Logrank p-value=0.0311). On contrary, patients with low expression of RNF43 substrate *VANGL1* (**D**.) had longer survival (Logrank test, p-value = 0.00518). Expression of DVL3, VANGL1 and ZNRF3 was analysed in the Figure 4 supplement 1A-D. **E**. Culture media from melanoma A375 WT and A375 IV cell lines were collected after 48 h and analyzed by Western blotting for presence of WNT5A. Densitometric analysis was done using the imageJ software. Equal number of cells was used. **F**. Schematic representation of genetic modification of A375 WT and A375 IV cells to stably overexpress exogenous RNF43 (+RNF43, grey) and to knockout *RNF43/ZNRF3* (R/Z dKO, dark grey) by CRISPR/Cas9 mediated gene editing. **G**. Effects of the RNF43 overexpression and *RNF43/ZNRF3* knockout in A375 WT and in its invasive derivate A375 IV. Exogenous RNF43 expression blocked DVL2 and DVL3 activation (arrowheads). Removal of endogenous RNF43 and ZNRF3 proteins presence had an opposite effect, N=6. Quantification is in the Figure 4 figure supplement 1E, F. Expression of the *WNT5A, RNF43, ZNRF3*, DVL2 and DVL2 in tested cell lines was checked and shown is in the Figure 4 figure supplement 2. **H. I**. Western blot showing DVL2 and DVL3 phosphorylation (arrowheads) in response to the 40 and 80 ng/ml 3h-long rhWNT5A treatments in A375 WT (H) and A375 IV (I) derived cell lines. β-ACTIN served as a loading control. LGK-974 was used to block endogenous Wnt ligands secretion and RNF43 was probed by HA antibody, N=3.

### RNF43 inhibits invasive properties of melanoma cells in vitro

A375 is a human melanoma cell line that is broadly used to study WNT5A role in melanoma (Anastas et al., 2014; Connacher et al., 2017; Da Forno et al., 2008; Ekström et al., 2014; Linnskog et al., 2016; Liu et al., 2018). For the purpose of our studies, we chose A375 wild type (WT) cells and their derivate with the increased metastatic potential referred to as A375 IV (Kucerova et al., 2014). Both A375 variants express *RNF43, WNT5A* (Figure 4 figure supplement 2A), and secrete WNT5A to the culture medium (Fig. 4E). Interestingly, *RNF43* expression in the A375 IV cells was significantly lower than in the A375 WT parental cells (Figure 4 figure supplement 2B). Expression of *ZNRF3* did not differ and it was not affected by RNF43 overexpression (Figure 4 figure supplement 2C). In order to study the RNF43 function, we generated A375 cells lacking *RNF43/ZNRF3* by CRISPR/Cas9 method (sequencing results are present in the Supplementary Table 1) and cells stably overexpressing RNF43 (Fig. 4F). The initial characterization of A375 derivatives essentially confirmed the findings from T-REx 293 (see Fig. 1) where RNF43 loss- and gain-of-function correlated strongly with the level of Wnt pathway activation assessed as DVL phosphorylation (Fig. 4G, quantified in Figure 4 figure supplement 1E, F). Total protein levels of DVL2, DVL3 as well as their expression remained unaffected by the manipulation of RNF43 expression (Figure 4 figure supplement 2D-G). Similarly to T-REx 293 cells, also in A375 WT (Fig. 4H) and A375 IV (Fig. 4I) melanoma cells RNF43 overexpression efficiently blocked WNT5A-induced signaling.

WNT5A signaling has been related to the numerous biological features that support invasive properties of melanoma (Arozarena and Wellbrock, 2017a; O’Connell and Weeraratna, 2009; Prasad et al., 2015; Weeraratna et al., 2002). To address if RNF43 affects any of these WNT5A-controlled properties, we have compared parental and RNF43-derivatives of A375 cells in the panel of functional assays that included: (i) wound healing assay, (ii) matrigel invasion assay, (iii) invadopodia formation assay and (iv) gelatin degradation assay. Firstly, cells overexpressing RNF43 showed suppressed 2D collective migration in the wound healing assay (Fig. 5A). Similarly, invasion of individual cells through the extracellular matrix (ECM) mimicking Matrigel was reduced by RNF43 (Fig. 5B). The analysis of invadopodia - specialized structures mediating adhesion and remodeling of surrounding ECM (Eddy et al., 2017; Masi et al., 2020), showed that cells overexpressing RNF43 formed less of them (Fig. 5C). In agreement, we also observed that these structures displayed reduced gelatin degradation activity in A375 WT and A375 IV cells overexpressing RNF43 (Fig. 5D). Further, treatment with WNT5A enhanced gelatin degradation capacity of A375 WT cells, but not their RNF43 overexpressing derivate (Fig. 5D). Representative images from conducted assays are shown in Figure 5 figure supplement 1-4. All these assays strongly support the conclusion that RNF43 acts as the strong molecular inhibitor of WNT5A-triggered pro-invasive features of melanoma.

**Figure 5.**
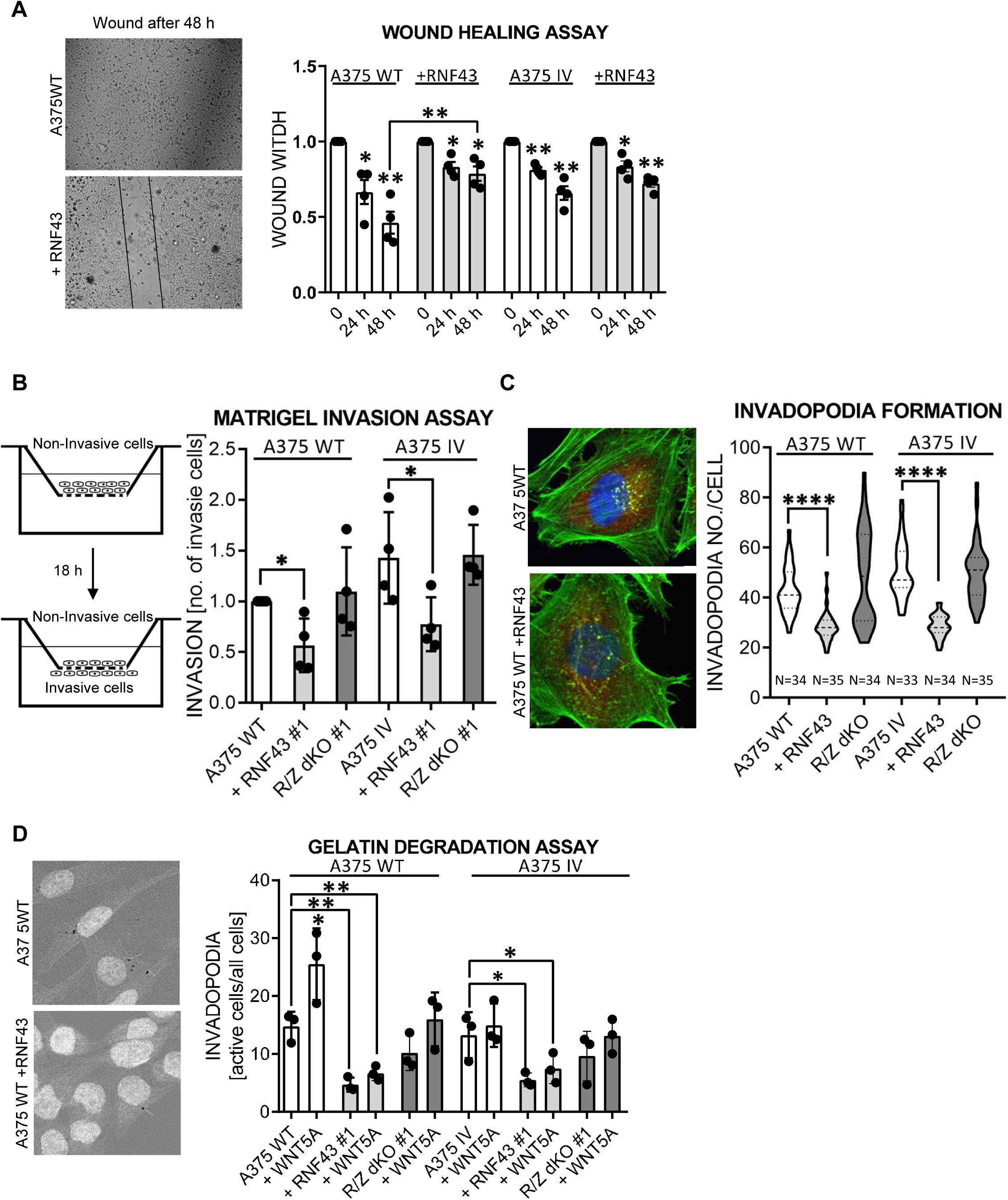
RNF43 inhibits WNT5A dependent invasive properties of human melanoma. **A**. RNF43 reduced melanoma cells migration in the wound healing assay. Wound width was tested at 24 h and 48 h after scratch, results were normalized to 1, as the scratch size at the experimental initial point. Cells proliferation was suppressed by serum starvation, unpaired two-tailed t-test: *p < 0.05, **p < 0.01, N=4. Representative photos at the end of the experiment are shown. **B**. Matrigel invasion assay – stable RNF43 overexpression inhibited invasive properties of the A375 WT and A375 IV. Serum starved cells were plated onto Matrigel coated porous membrane. Medium containing 20% of serum was used as chemoattractant. After 18 h of incubation cells were fixed in methanol, noninvaded ones were removed from the upper part of transwell insert by cotton swab. Results were normalized to 1 for the number of invaded A375 WT cells, unpaired two-tailed t-test: *p < 0.05, N=4. Representative photos are shown in the Figure 5 figure supplement 1. **C**. Quantification of the invadopodia formed by melanoma cells. RNF43 overexpression in the A375 WT and A375 IV decreased number of invadopodia, based on the analysis of confocal images. Number of cortactin/F-actin double positive puncta in the individual cells was calculated in the imageJ software, unpaired two-tailed t-test: ****p<0.0001. Examples of confocal imaging are shown: green – phalloidin, red – cortactin, blue – DNA. See Figure 5 figure supplement 2 for images from all experimental conditions. **D**. Gelatin degradation assay-both A375 WT and A375 IV RNF43-overexpressing cell lines showed decreased capacity to locally degrade the extracellular matrix modification. Serum starved cells were plated onto gelatin-Oregon Green coated coverslips and incubated for 24 hours. Images obtained by Leica SP8 confocal microscope were analyzed for the presence of gelatin degradation by individual cells using imageJ software, unpaired two-tailed t-test: *p < 0.05, **p < 0.01, N=3. Example of gelatin degradation is shown, more pictures are present in the Figure 5 figure supplement 3 and 4. Data used in the A – B is present in the Figure 5 Data Source file.

### RNF43 prevents acquisition of resistance to BRAF V600E targeted therapy

The Mitogen activated protein kinase (MAPK) pathway is hyperactivated in melanoma (Davies et al., 2002) as a result of UV-induced mutations triggering constitutive activation of this signaling axis. The most common genetic aberration - *BRAF V600E* is a target of anti-melanoma therapy (Akbani et al., 2015; Birkeland et al., 2018; Chapman et al., 2011; Flaherty et al., 2010; Hodis et al., 2012; Shain et al., 2015). Drugs targeting mutated BRAF (e.g. Vemurafenib/PLX4032) in melanoma improved patient’s survival (Chapman et al., 2011; Flaherty et al., 2010; Joseph et al., 2010). Unfortunately, patients receiving BRAF inhibitors (BRAFi) relapses after several months of monotherapy because of the acquired resistance (Nazarian et al., 2010). WNT5A was shown to play a crucial role in the process leading to the Vemurafenib resistance (Anastas et al., 2014; Mohapatra et al., 2018; O’Connell et al., 2013; Prasad et al., 2015; Webster et al., 2015). Therefore, we were interested to check whether RNF43 inhibits via its effects on WNT5A signaling cellular plasticity in response to Vemurafenib (PLX4032), a clinically used *BRAF* V600E inhibitor.

The process of Vemurafenib resistance acquisition can be modelled in vitro. We applied experimental scheme optimized for A375 (Anastas et al., 2014). This model (Fig. 6A) allows to study both acute responses to Vemurafenib (24 h treatment) as well as the gradual adaptation the long-term cell culture in the increasing vemurafenib doses. Vemurafenib resistant (VR) cells can be obtained after approximately 2 months. As shown in Fig. 6B, treatment with Vemurafenib resulted in rapid and complete inhibition of ERK1/2 phosphorylation, the readout of MAPK activation (compare lane 1 and 2). In contrast A375 WT VR cells showed constitutive ERK1/2 phosphorylation even in the presence of 2 μM Vemurafenib (compare lane 2 and 3). Interestingly, transient exposition to Vemurafenib resulted in the impeded phosphorylation of ROR1, DVL2 and DVL3 (Fig. 6B, C, D). On the other side, VR cells displayed elevated ROR1 levels and increased phosphorylation of DVL2 and DVL3 (Fig. 6B, C and D). This suggests that activation of the noncanonical WNT5A-induced signaling is indeed a part of the melanoma adaptation to Vemurafenib.

**Figure 6.**
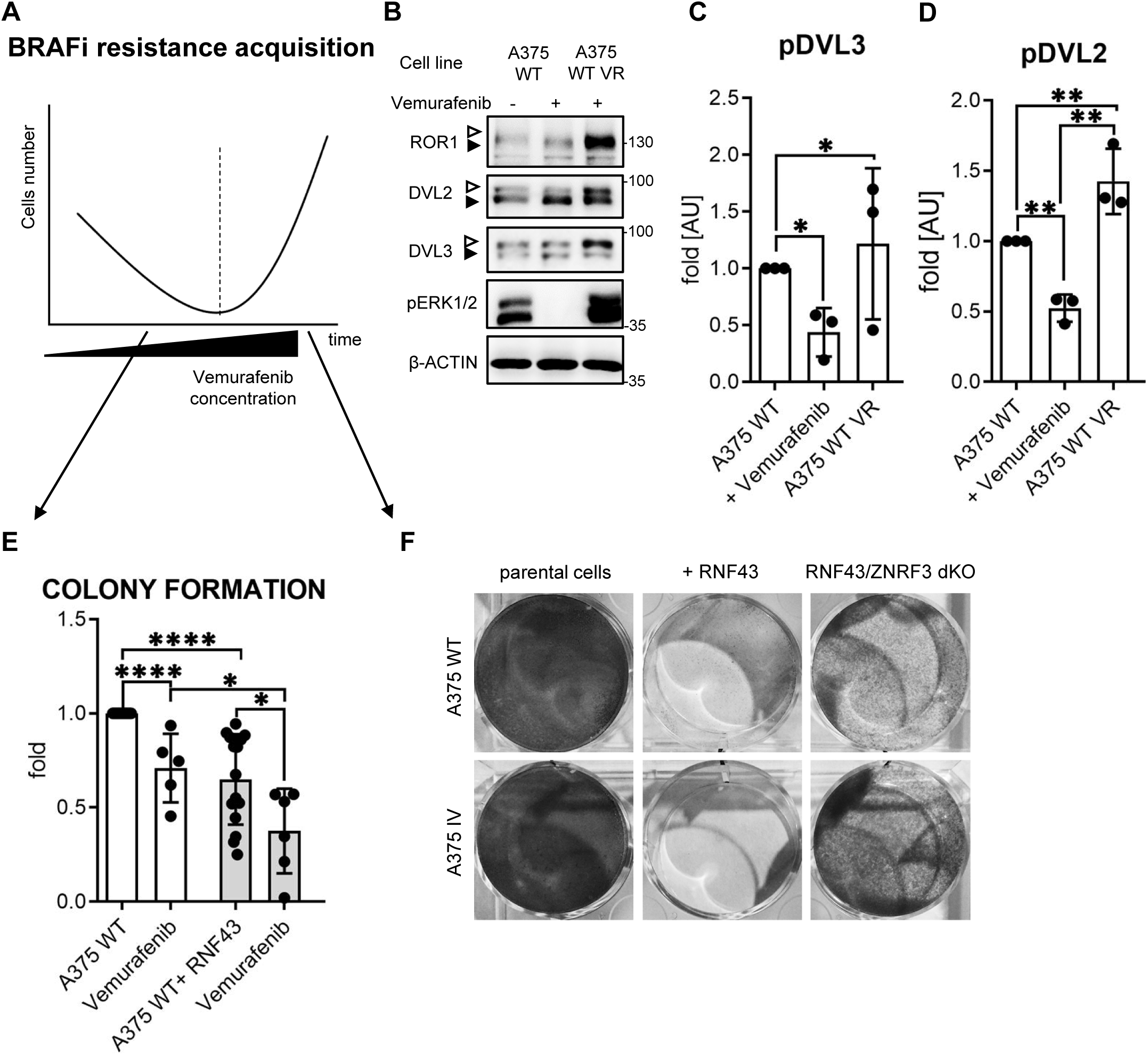
RNF43 overexpressing melanoma cells do not develop resistance to bRAF inhibition. **A**. Scheme showing the experimental model used for the analysis of vemurafenib resistance acquisition. Melanoma cells are exposed to the increasing doses of the *BRAF* V600E inhibitor Vemurafenib and following initial decrease in cell numbers recover and obtain capacity to grow in the presence of Vemurafenib. **B**., **C**., **D**. Western blot analysis of the cellular responses to the acute Vemurafenib treatment (0.5 µM, 24 hours) in comparison to the signaling in Vemurafenib-resistant (VR) cells growing in presence of 2 µM Vemurafenib. In VR cells ERK1/2 is constitutively phosphorylated even in the Vemurafenib presence. β-ACTIN served as a loading control. A375 WT VR cells showed increased activation of DVL2 and DVL3 (arrowheads: DVL2, DVL3, quantifications in C and D) and higher expression of ROR1. Unpaired two-tailed t-test: *p < 0.05, **p < 0.01, N=3. **E**. Melanoma cell lines A375 WT and A375 IV overexpressing RNF43 showed decreased ability to grow and form colonies when seeded in the low density. Colonies were fixed and stained with crystal violet after seven days. Paired (Vemurafenib – vs +) and unpaired (WT vs IV) two-tailed t-tests: *p < 0.05, ****p < 0.0001, N ≥ 5. **F**. RNF43 overexpressing A375 WT and A375 IV did not develop resistance to the BRAF V600E inhibition by vemurafenib treatment. Cells were cultured for approximately two months in the presence of increasing doses of the inhibitor. Photos show crystal violet stained cultures at the end of the selection process. Data used in the C, D and E is shown in the Figure 6 Source data file.

Therefore, we challenged with Vemurafenib A375 WT and its RNF43 expressing derivatives. As shown in Fig. 6E exogenous RNF43 decreased colony formation and proliferation of cells seeded in the low density and Vemurafenib further enhanced this effect. Importantly, both A375 WT and A375 IV overexpressing RNF43 completely failed to develop resistance to Vemurafenib and died off during the selection at 1 μM Vemurafenib concentration (Fig. 6F). Altogether these data confirm earlier findings on the importance of WNT5A signaling in the acquisition of Vemurafenib resistance and demonstrate that RNF43 can completely block this process.

## Discussion

Our study identified RNF43 as the inhibitor of noncanonical WNT5A-induced signaling. RNF43 physically interacted with multiple receptor components of the Wnt/PCP pathway such as ROR1/2, VANGL1/2 or DVL1/2/3 and triggered degradation of VANGL2 and membrane clearance of ROR1; ultimately resulting in the reduced cell sensitivity to WNT5A. The newly discovered RNF43 action in WNT5A-mediated signaling seems to be mechanistically different than the well-known function in the Wnt/β-catenin pathway. For example, we observed ROR1 and VANGL2 interaction with RNF43 in the absence of DVL. In contrast, DVL seems to be essential for the activity of RNF43 in the Wnt/β-catenin pathway (Jiang et al., 2015). Further, inhibitory action of RNF43 in WNT5A-signaling could not be blocked by inhibition lysosomal pathway, in contrast to the earlier observations in WNT/β-catenin pathway (Koo et al., 2012). On the other side, WNT5A signaling can be similarly to Wnt/β-catenin promoted by RNF43 inhibitors from R-SPO family. Also, in line with the earlier findings that RNF43 leads to the packing of ubiquitinated FZD to the RAB5^+^ endosomes (Koo et al., 2012), ROR1 is as well internalized via clathrin-dependent mechanism into RAB5^+^ endosomes. It remains to be studied how RNF43 in a coordinated manner controls both WNT/b-catenin and noncanonical WNT pathways.

We demonstrate that the newly characterized RNF43-WNT5A regulatory module controls WNT5A signaling and biology in melanoma. WNT5A-induced signaling plays in melanoma a crucial role. Up to date, 5-year survival of metastatic melanoma patients rate between 5-19%, depending by the location and the number of metastases (Sandru et al., 2014). Elevated expression of *WNT5A*, associates with negative overall survival in melanoma (Da Forno et al., 2008; Luo et al., 2020; Weeraratna et al., 2002) - we have observed inverse correlation for *RNF43*, which was a positive prognostic factor in melanoma and got silenced as melanoma progressed. WNT5A promotes multiple pro-invasive features of melanoma cells such as EMT, invasion, metastasis, cell proliferation and extracellular matrix remodeling by melanoma cells (Dissanayake et al., 2008, 2007; Fernández et al., 2016; Lai et al., 2012). RNF43 overexpression efficiently suppressed all tested pro-metastatic properties of melanoma cells associated with WNT5A. Among those, the clinically most relevant is the acquisition of resistance to BRAF inhibitor Vemurafenib.

*BRAF V600E* mutation appears in up to 50% of melanoma cases results in the oncogenic activation of MAPK pathway (Akbani et al., 2015; Wan et al., 2004). Vemurafenib (PLX4032), a compound selectively inhibiting BRAF V600E, showed positive clinical effects in melanoma (Bollag et al., 2012; Joseph et al., 2010). Unfortunately, most of the patients develop resistance to Vemurafenib treatment and progress (Chapman et al., 2011). Multiple mechanisms underlying acquisition of resistance were described (Arozarena and Wellbrock, 2019, 2017b; Johnson et al., 2015; Luebker and Koepsell, 2019; Schmitt et al., 2019; Su et al., 2020, 2017; Talebi et al., 2018; Tirosh et al., 2016). Among those mechanisms, WNT5A signaling has a prominent role - *WNT5A* expression was shown to positively correlate with Vemurafenib resistance (Anastas et al., 2014; Prasad et al., 2015; Webster et al., 2015) and WNT5A treatment decreased melanoma cells response to the Vemurafenib (Anastas et al., 2014; O’Connell et al., 2013). Our finding that RNF43-controlled regulatory axis could completely block development of resistance to BRAF inhibition further highlights importance of WNT5A signaling in this process and also uncovers a mechanism that can be explored therapeutically.

Relevance of our findings is likely not limited to melanoma. Signaling cascade RSPO– LGR4/5–RNRF43/ZNRF3 has been shown to regulate variety of biological processes. In light of our results, it is tempting to speculate that WNT5A-RNF43 axis regulates other developmental, physiological and patho-physiological conditions. For example, *WNT5A* is overexpressed in gastric cancer where it positively correlates with the presence of the lymph node metastasis, tumor depth, EMT induction and poor prognosis (Astudillo, 2020; Hanaki et al., 2012; Kanzawa et al., 2013; Kurayoshi et al., 2006; Nam et al., 2017; Saitoh et al., 2002). Notably, reduced RNF43 function is a negative prognosis factor in gastric cancer patients (Gao et al., 2017; Neumeyer et al., 2019a; Niu et al., 2015) and RNF43 loss of function type of mutation exacerbated *Helicobacter pylori*-induced gastric tumor carcinogenesis associated with the upregulation of *WNT5A* mRNA level (Katoh, 2007; Li et al., 2014; Neumeyer et al., 2019b; Peek and Crabtree, 2006). Further, in colorectal cancer RNF43 mutations were found to associate with *BRAF* V600E mutation (Matsumoto et al., 2020; Yan et al., 2017).These results suggest the existence of more universal functional WNT5A-RNF43 axis where RNF43 acts as a gatekeeper guarding the abnormal pro-cancerogenic noncanonical Wnt pathway activation.

Further exciting avenues relate to the importance of RSPO-RNF43/ZNRF3 module in the regulation of multiple developmental processes dependent on WNT5A. There are literature hints that suggest that indeed WNT5A-signaling is fine-tuned by RNF43/ZNRF3 during convergent extension movements. The regulation of Rspo3 has been proven in *Xenopus* embryogenesis, where it regulates gastrulation movements and head cartilage morphogenesis in a manner involving Wnt5a and Syndecan-4 binding by R-spondin. Strikingly, *Rspo3* antisense morpholino caused phenotype characteristic for the noncanonical Wnt signalling pathway - *spina bifida* (Ohkawara et al., 2011). Similarly, overexpression of *Znrf3* in zebrafish embryo caused shortened body axis and abnormal shape of somites, phenotypes also recognised as typical for Wnt/PCP pathway perturbances (Hao et al., 2012). And, finally in mammals, a fraction of *Znrf3* KO mice showed an open neural tube phenotype (Hao et al., 2012), again reminiscent of defective Wnt/PCP signalling. Altogether, these observations together with our data suggest that RSPO-RNF43/ZNRF3 signaling represents an evolutionary conserved and widely used mechanism used to control activation of noncanonical WNT signaling.

## Materials and Methods

### 1. Cell lines and treatments

T-REx™-293 (R71007, Thermo Fisher Scientific), GFP labelled human melanoma A375 wild type (WT) and its metastatic derivate A375 IV cell lines (Kucerova et al., 2014) were propagated in the Dulbecco’s modified Eagle’s medium (DMEM, 41966–029, Gibco, Life Technologies) supplemented with the 10% fetal bovine serum (FBS, 10270–106, Gibco, Life Technologies), 2 mM L-glutamine (25030024, Life Technologies), 1% penicillin-streptomycin (XC-A4122/100, Biosera) under 5% (vol/vol) CO2 controlled atmosphere at 37 °C.. For inhibition of endogenous the Wnt ligands, cells were treated with the 0.5 μM Porcupine inhibitors C-59 (ab142216, Abcam) or LGK-974 (1241454, PeproTech). For canonical Wnt signaling activation recombinant the human WNT3A (CF 5036-WN-CF, RnD Systems) was used and the recombinant human WNT5A (645-WN-010, RnD Systems) for noncanonical Wnt pathway stimulation, both in 40 ng/ml, 60 ng/ml or 80 ng/ml concentrations for 3h or overnight treatments. Co-treatment with the recombinant human R-Sponidin-1 (120-38, PeproTech) in 50 ng/ml dose was applied where indicated. Dansylcadaverine (D4008, Sigma-Aldrich) 50 µM treatment along with 3 h tetracycline was applied to block clathrin dependent endocytosis pathway (Blitzer and Nusse, 2006).

For preparation of stable cell lines, antibiotic selection after plasmid DNA transfection was performed using 5 μg/ml blasticidin S (3513-03-9, Santa Cruz Biotechnology) or 200 μg/ml of hygromycin B (31282-04-9, Santa Cruz Biotechnology) for T-REx-293 cells and accordingly 400 μg/ml and 5 μg/ml in case of A375 melanoma cell line. As a result, tetracycline inducible T-REx-293 RNF43 and RNF43 Mut1 TetON, T-REx-293 *RNF43*/*ZNRF3* dKO RNF43 TetON, A375 WT +RNF43 and A375 IV +RNF43 were obtained. T-REx-293 *DVL1*/*2*/*3* tKO cells were described previously (Paclíková et al., 2017). For transgene expression induction (TetON), T-REx-293 cells were treated with the 1 μg/ml of tetracycline (60-54-8, Santa Cruz Biotechnology) for indicated time (3 hours to overnight). Lysosomal degradation pathway was blocked by the 10 μM Chloroquine (C662, Sigma) treatment, whereas 10 μM MG-132 (C2211, Sigma) was used for the proteasome inhibition. Generation of the melanoma cells resistant to Vemurafenib (HY-12057, MedChem Express) was performed according to the published protocols, i.e. (Anastas et al., 2014). Resistant cells were cultured in the presence of 2 μM vemurafenib. For transient treatments (24h or 48 h) of melanoma cell lines, 0.5 μM vemurafenib has been used.

### 2. Plasmids/cloning

Backbone of the plasmid pcDNA4-TO-RNF43-2xHA-2xFLAG (kindly gifted by Bon-Kyoung Koo together with pcDNA4-TO-RNF43Mut1-2xHA-2xFLAG (Koo et al., 2012)) was used for further cloning. Briefly, for generation of the BioID inducible pcDNA4-TO-RNF43-BirA*-HA plasmid, cDNA encoding RNF43 without stop codon was amplified by the PCR and cloned into the pcDNA3.1 MCS-BirA(R118G)-HA (Addgene plasmid #36047) using HpaI (ER1031, Thermo Fisher Scientific) and EcoRI (ER0271, Thermo Fisher Scientific) restriction enzymes to fuse it in frame with the BirA*-HA sequence. Then, RNF43-BirA*-HA cDNA was amplified and cloned by the In-Fusion cloning method (639690, Takara Bio) into linearized by HindIII (ER0501, Thermo Fisher Scientific) and XbaI (ER0681, Thermo Fisher Scientific) pcDNA4-TO plasmid. To eliminate BirA* enzyme mediated potential false positive results, pcDNA3-RNF43-HA was prepared by subcloning RNF43 PCR product containing HA encoding sequence in reverse primer to the pcDNA3 backbone (Invitrogen). All obtained plasmids were verified by the Sanger sequencing method.

Other plasmids used were described previously and included: myc-Vangl1, GFP-Vangl2, GFP-Vangl2ΔN, GFP-Vangl2ΔC, GFP-Vangl2ΔNΔC (Belotti et al., 2012), pEGFP-C1-Rab5a (Chen et al., 2009), GFP-rab11 WT (Addgene #12674), His-Ubiquitin (Tauriello et al., 2010), pcDNA3-Flag-mDvl1 (Tauriello et al., 2010), pCMV5-3xFlag Dvl2 (Addgene #24802), pCDNA3.1-Flag-hDvl3 (Angers et al., 2006), pcDNA3.1-hROR1-V5-His (gifted by Kateřina Tmějová), pcDNA3-Ror2-Flag and pcDNA3-Ror2-dCRD-FLAG (Sammar et al., 2004), pRRL2_ROR1ΔCYTO and pRRL2_ROR1ΔTail (Gentile et al., 2011), hCas9 (Addgene #41815), gRNA_GFP-T1 (Addgene #41819), PiggyBack-Hygro and Transposase coding plasmids (gifted by Bon-Kyoung Koo). Sequences of primers used for cloning is present in the Table 1.

**Table 1.**
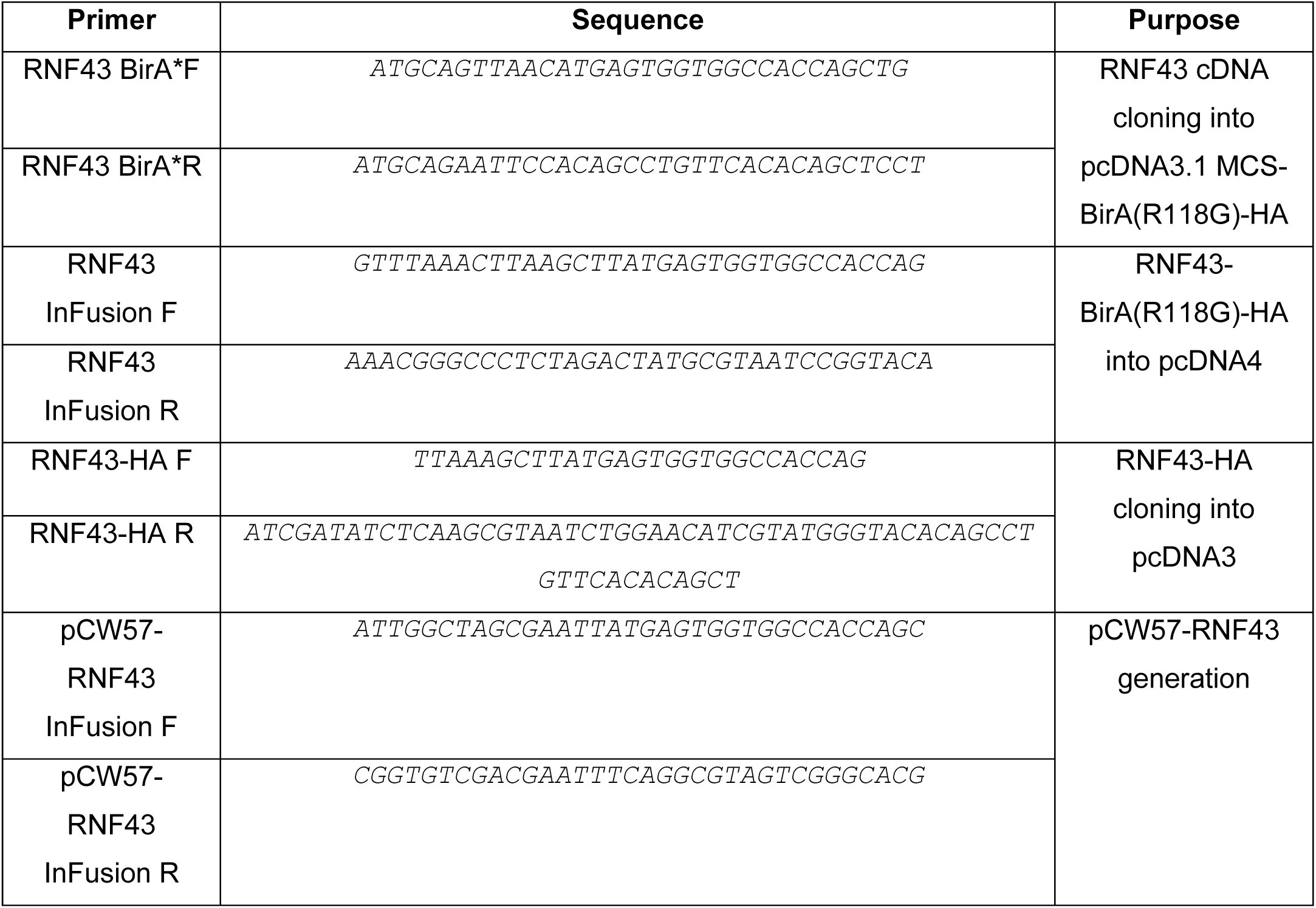
**Cloning and mutagenesis primers**

### 3. CRISPR/Cas9

For targeting *RNF43* and *ZNRF3* in the T-Rex-293, gRNAs: *TGAGTTCCATCGTAACTGTG<u>TGG</u>* (<u>PAM</u>) and *AGACCCGCTCAAGAGGCCGG<u>TGG</u>* were cloned into gRNA_GFP-T1 backbone and transfected together with PiggyBack-Hygro and Transposase coding plasmids using polyethylenimine (PEI) in a way described below. For *ROR1* and *WNT5A* knock-out cell lines generation, gRNA *CCATCTATGGCTCTCGGCTG<u>CGG</u>* (ROR1) and *AGTATCAATTCCGACATCGA<u>AGG</u>* (WNT5A) were used. Transfected cells were hygromycine B selected and seeded as single cells. Genomic DNA isolation was performed using DirectPCR Lysis Reagent (Cell) (Viagen Biotech), Proteinase K (EO0491, Thermo Fisher Scientific) and DreamTaq DNA Polymerase (EP0701, Thermo Fisher Scientific) according to the manufacturers. PCR products were analyzed by restriction digestion using Taal (ER1361, Thermo Fisher Scientific) in case of *RNF43*, HpaII (ER0511, Thermo Fisher Scientific) – *ZNRF3*, TaqI (ER0671, Thermo Fisher Scientific) - *WNT5A* and TseI (R0591S, New England BioLabs) - *ROR1* for detection of Cas9 mediated disruptions in the recognition sites.

For targeting *RNF43*/*ZNRF3* in the A375 and in the A375 IV melanoma lines, gRNAs: *AGTTACGATGGAACTCA<u>TGG</u>* (RNF43) and *CTCCAGACAGATGGCACAGT<u>CGG</u>* (ZNRF3) were accordingly cloned by described protocol into the pU6-(BbsI)CBh-Cas9-T2A-mCherry (Addgene #64324) and pSpCas9(BB)-2A-GFP (PX458) (Addgene #48138) backbones, transfected and sorted as single, GFP and mCherry double positive cells. Then analyzed by restriction enzymes Hin1II (ER1831, Thermo Fisher Scientific) and Taal as described above. Finally, PCR products were sequenced using the Illumina platform and compared with the reference sequence (Malcikova et al., 2015). Sequencing results are presented in the Supplementary Table 1.

### 4. RNF43 BioID analysis

Following IP washes, bead bound protein complexes were processed directly on beads covering protein reduction (50mM dithiothreitol – DTT, 30min, 37°C), alkylation (50mM iodacetamide – IAA, 30min, 25°C, dark; IAA excess quenched by additional DTT) and trypsin digestion (750ng of sequencing grade trypsin, Promega) in 50mM NaHCO_3_ buffer. Beads were incubated at 37°C with mild agitation for 14 hours. Resulting peptides were extracted into LC-MS vials by 2.5% formic acid (FA) in 50% acetonitrile (ACN) and 100% ACN with addition of polyethylene glycol (20,000; final concentration 0.001%) (Stejskal et al., 2013) and concentrated in a SpeedVac concentrator (Thermo Fisher Scientific).

LC-MS/MS analyses of all peptide mixtures were done using RSLCnano system (SRD-3400, NCS-3500RS CAP, WPS-3000 TPL RS) connected to Orbitrap Elite hybrid spectrometer (Thermo Fisher Scientific). Prior to LC separation, tryptic digests were online concentrated and desalted using trapping column (100 μm × 30 mm, 40°C) filled with 3.5-μm X-Bridge BEH 130 C18 sorbent (Waters). After washing of trapping column with 0.1% FA, the peptides were eluted (flow 300 nl/min) from the trapping column onto an analytical column (Acclaim Pepmap100 C18, 3 µm particles, 75 μm × 500 mm, 40°C; Thermo Fisher Scientific) by 100 min nonlinear gradient program (1-56% of mobile phase B; mobile phase A: 0.1% FA in water; mobile phase B: 0.1% FA in 80% ACN). Equilibration of the trapping column and the column was done prior to sample injection to sample loop. The analytical column outlet was directly connected to the Digital PicoView 550 (New Objective) ion source with sheath gas option and SilicaTip emitter (New Objective; FS360-20-15-N-20-C12) utilization. ABIRD (Active Background Ion Reduction Device, ESI Source Solutions) was installed.

MS data were acquired in a data-dependent strategy selecting up to top 10 precursors based on precursor abundance in the survey scan (350-2000 m/z). The resolution of the survey scan was 60 000 (400 m/z) with a target value of 1×10^6^ ions, one microscan and maximum injection time of 200 ms. HCD MS/MS (32% relative fragmentation energy) spectra were acquired with a target value of 50 000 and resolution of 15 000 (400 m/z). The maximum injection time for MS/MS was 500 ms. Dynamic exclusion was enabled for 45 s after one MS/MS spectra acquisition and early expiration was disabled. The isolation window for MS/MS fragmentation was set to 2 m/z.

Data are available via ProteomeXchange (Deutsch et al., 2020) with identifier PXD020478 in the PRIDE database (Perez-Riverol et al., 2019). The analysis of the mass spectrometric RAW data files was carried out using the MaxQuant software (version 1.6.2.10) using default settings unless otherwise noted. MS/MS ion searches were done against modified cRAP database (based on http://www.thegpm.org/crap) containing protein contaminants like keratin, trypsin etc., and UniProtKB protein database for *Homo sapiens* (ftp://ftp.uniprot.org/pub/databases/uniprot/current_release/knowledgebase/reference_proteomes/Eukaryota/UP000005640_9606.fasta.gz; downloaded 19.8.2018, version 2018/08, number of protein sequences 21,053). Oxidation of methionine and proline, deamidation (N, Q) and acetylation (protein N-terminus) as optional modification, carbamidomethylation (C) as fixed modification and trypsin/P enzyme with 2 allowed miss cleavages were set. Peptides and proteins with FDR threshold <0.01 and proteins having at least one unique or razor peptide were considered only. Match between runs was set among all analyzed samples. Protein abundance was assessed using protein intensities calculated by MaxQuant. Protein intensities reported in proteinGroups.txt file (output of MaxQuant) were further processed using the software container environment (https://github.com/OmicsWorkflows), version 3.7.2a. Processing workflow is available upon request. Briefly, it covered: a) removal of decoy hits and contaminant protein groups, b) protein group intensities log2 transformation, c) LoessF normalization, d) imputation by the global minimum and e) differential expression using LIMMA statistical test. Prior to volcano plot plotting, suspected BirA* binders were filtered out (proteins identified on at least 2 peptides in both technical replicates of particular BirA* sample, and present in >3 samples). Volcano plot was created in R using ggplot2 and ggrepel R packages by R version 3.6.1. Proteins with adjusted p-value <0.05 and log fold change >1 were further subjected to gene ontology tools, considering only the first ID of majority protein IDs: g:Profiler online tool (https://biit.cs.ut.ee/gprofiler/gost,version e98_eg45_p14_ce5b097) (Raudvere et al., 2019) was used and selected GO terms were highlighted. RNF43 interactors from BioID assay are listed in the Figure 1 Supplementary table 1 and results obtained by g:Profiler are present in the Figure 1 Supplementary table 2.

### 5. Transfection

T-REx™-293 cells were transected using 1 μg/ml, pH 7.4 polyethylenimine (PEI) and plasmid DNA in 4:1 ratio (Paclíková et al., 2017). Plasmid DNA in amount of 3 µg for 6 cm culture dish (ubiquitination assay) and 6 µg for 10 cm dish (co-immunoprecipitation or stable cell lines preparation). Approximately 1×10^6^ of A375 and A375 IV cells were electroporated with 6 μg of plasmid DNA utilizing Neon Transfection System (Thermo Fisher Scientific) 1200V, 40 ms, 1 pulse. Culture media were changed six hours post-transfection.

### 6. His-ubiquitin pulldown assay

Cells were transfected with the plasmid encoding polyhistidine-tagged ubiquitin, RNF43-HA or enzymatically inactive RNF43, protein of interest and cultured overnight. Next, cells were treated with 0.2 µM epoxomicin (E3652, Sigma) for 4 hours and lysed in the buffer containing: 6M guanidine hydrochloride (G3272, Sigma), 0.1 M Na_x_H_x_PO_4_ pH 8.0 and 10 mM imidazole (I5513, Sigma), sonicated and boiled. Insoluble fraction was removed by the centrifugation (16 000g, RT, 10 min). For the pull down of tagged proteins, 10 µl of equilibrated in lysis buffer His Mag Sepharose beads Ni (GE28-9799-17, GE Healthcare) was added to each sample and kept on a roller overnight. Then, beads were washed three times in the buffer containing 8M urea (U5378, Sigma),0.1 M Na_x_H_x_PO_4_ pH 6.3, 0.01 M Tris and 15 mM imidazole, resuspended in the 100 μl of Western blot sample buffer, boiled for 5 minutes and loaded onto SDS-PAGE gel. Approximately 10% of cellular lysate was used as a transfection control after ethanol precipitation and resuspension in the Western blot sample buffer.

### 7. Western blotting and antibodies

Western blot analysis was performed as it was described before using samples with same protein amount, measured by the DC Protein Assay (5000111, Bio-Rad), or lysed directly in the sample buffer (2% SDS, 10% glycerol, 5% β-mercaptoethanol, 0.002% bromphenol blue and 0.06 M Tris HCl, pH 6.8 and Protease inhibitor cocktail (11836145001,Roche) after PBS wash (Mentink et al., 2018). Protein extraction from mouse tissues was done by homogenizing in the 1% SDS, 100 mM NaCl, 100 mM Tris, pH 7.4 buffer, sonication, clarification by centrifugation (16000g. 4°C, 15 min) and protein concentration measurement. Next, 25 μg of protein samples was mixed with Western blot sampling buffer and loaded onto SDS-PAGE gels. Briefly, after electrophoretic separation, proteins were transferred onto Immobilon-P PVDF Membrane (IPVH00010, Millipore) and detected using primary and corresponding HRP-conjugated secondary antibodies on Fusion SL imaging system (Vibler) using Immobilon Western Chemiluminescent HRP Substrate (Merck, WBKLS0500). Molecular size of bands is marked in each panel [kDa]. List of used antibodies is present in the Table 2.

**Table 2.**
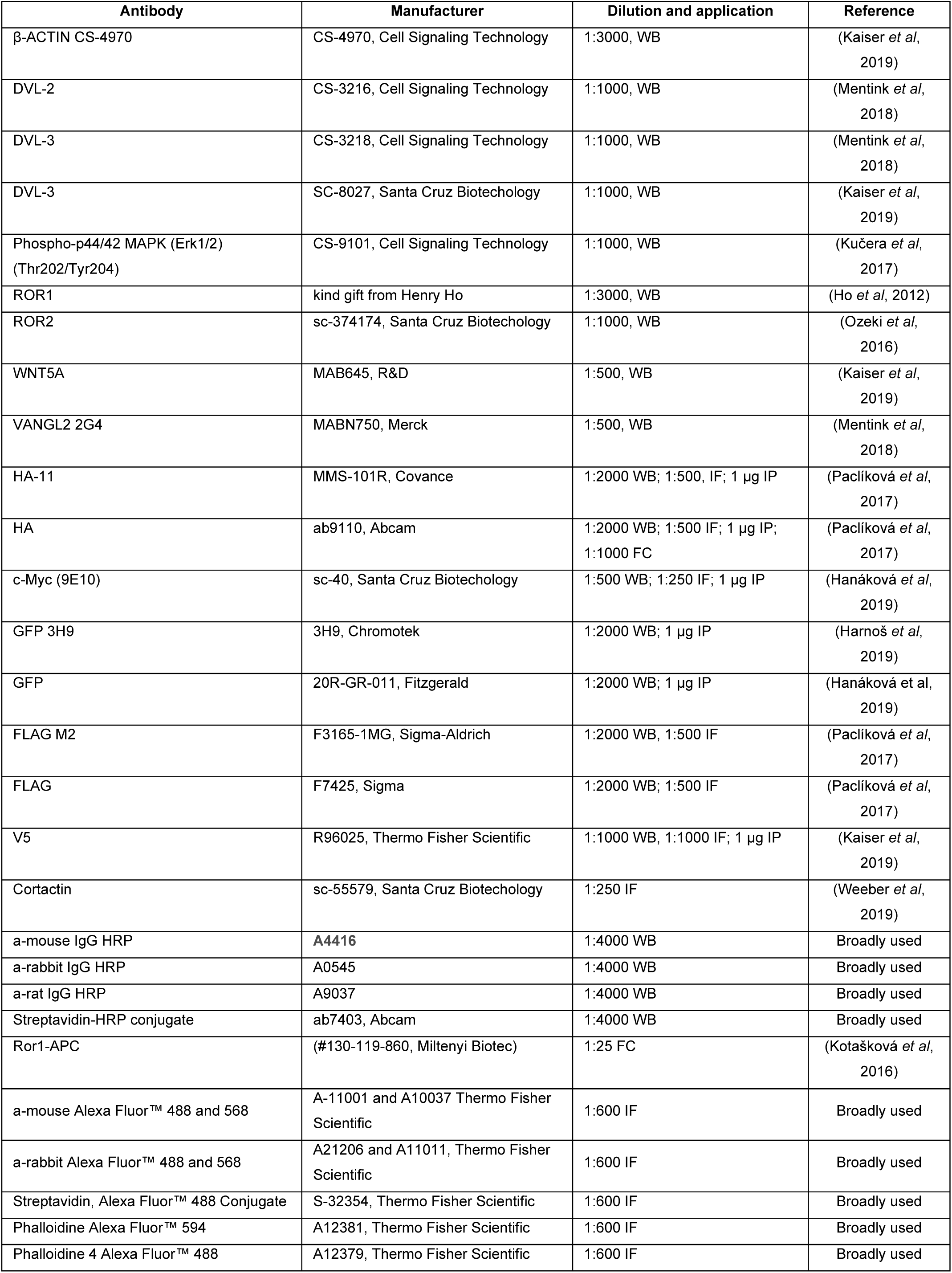
**Antibodies**

### 8. Immunofluorescence and confocal microscopy

Cells growing on the glass were fixed in 4% paraformaldehyde (PFA) in PBS. Fixed cells were permeabilized by the 0.1% Triton X-100 in PBS and blocked in the 1% solution of bovine serum albumin in PBS. Then, samples were incubated overnight at 4°C with primary antibodies diluted in the 1% BSA in PBS and washed. Corresponding Alexa Fluor secondary antibodies (Invitrogen) were incubated with samples for 1h at room temperature, along with the 1 µg/ml Hoechst 33342 (H1399, Thermo Fisher Scientific) for nuclei staining. After PBS washes, samples were mounted in the DAKO mounting medium (S3023, DAKO). Images were taken on the confocal laser scanning microscopy platform Leica TCS SP8 (Leica). For co-localization analysis histograms for each channel were prepared in the LAS X Life Science (Leica) software and plotted in the GraphPad Prism 8.

### 9. Immunoprecipitation

T-REx-293 cells were transfected with the proper plasmid DNA and cultured for 24 hours. Then, cells were washed two times with PBS and lysed for 15 min in the buffer containing 50 mM Tris pH7.6, 200 mM NaCl, 1 mM EDTA, 0.5% NP40, fresh 0.1mM DTT (E3876, Sigma) and protease inhibitor cocktail (04693159001, Roche). Insoluble fraction was removed by the centrifugation (16 000g, RT, 15 min), 10% of total cell lysate was kept as Western blot control. Lysates were incubated with the 1 μg of antibody for 16 h at 4°C on the head-over-tail rotator. Next, 20 μl of protein G-Sepharose beads (17-0618-05; GE Healthcare) equilibrated in the complete lysis buffer were added to each sample and incubated for 4 hours at 4°C, following six washes using lysis buffer and resuspension in 100 μl of Western blot sample buffer. Immunoprecipitation experiments were analyzed by the Western blot.

### 10. Flow cytometric determination of ROR1 surface expression

Determination of the ROR1 surface expression of T-REx™-293 and its derivates was performed using the anti-ROR1-APC (#130-119-860, Miltenyi Biotec) and Accuri C6 (BD Biosciences) (*RNF43*/*ZNRF3* dKO cells) or using BD FACSVerse Flow Cytometer (BD Biosciences) (TetON cells). Cells were harvested in 0.5 mM EDTA/PBS, washed in PBS and incubated in 2% FBS in PBS with anti-ROR1-APC antibody (1:25, #130-119-860, Miltenyi Biotec) on ice for 30 minutes. The cells were washed and resuspended in PBS, incubated with propidium iodide (10 ng/ml, #81845, Sigma-Aldrich) for 5 minutes to exclude dead cells from analysis. For the detection of ROR1 surface expression in HA positive cells, ROR1-APC stained cells were washed in PBS, fixed in 4% PFA at RT for 15 minutes, permeabilized in 0,02% Triton X-100 at RT for 15 minutes and incubated with anti-HA antibody (1:1000, #9110, Abcam) in staining buffer at RT for 30 minutes. After two washes, cells were incubated with secondary antibody ALEXA Fluor® 488 Donkey anti-Rabbit (#A21206, Invitrogen) at RT for 20 minutes, washed and measured using FACS Verse (BD Biosciences). Data were analyzed using NovoExpress® Software (ACEA Biosciences).

### 11. qPCR - quantitative polymerase chain reaction

Messenger RNA was isolated using the RNeasy Mini Kit (74106; Qiagen) according to the manufacturer’s instructions. One microgram of mRNA was transcribed to cDNA by the RevertAid Reverse Transcriptase (EP0442, Thermo Fisher Scientific) and analyzed by use of the LightCycler® 480 SYBR Green I Master (04887352001, Roche) and the Light Cycler LC480 (Roche). Results are presented as 2^−ΔΔ*CT*^ and compared by unpaired Student’s *t* test. Mean expression of *B2M* and *GAPDH* was used as reference. Primers are listed in the Table 3.

**Table 3.**
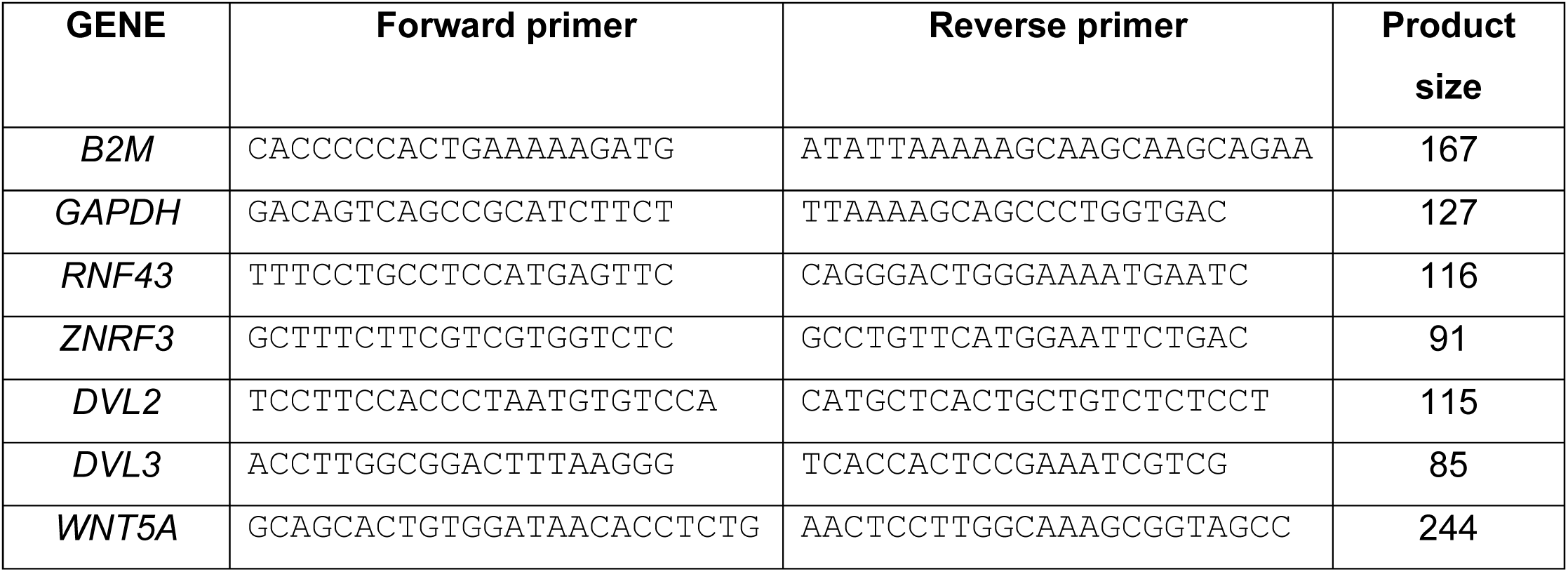
**qPCR primers**

## 12. Databases

RNF43, VANGL1, DVL3 genes expression on different melanoma stages was analyzed through the Oncomine (Rhodes et al., 2004) database in the different datasets (Talantov et al., 2005), (Xu et al., 2008), (Haqq et al., 2005). OncoLnc (Anaya, 2016) database was employed to elucidate whether expression of the *RNF43, ZNRF3, VANGL1* and *DVL3* genes expression have significant impact on the melanoma patients overall survival. RNF43 BioID data are available via ProteomeXchange (Deutsch et al, 2020) in the PRIDE database (PXD020478) (Perez-Riverol et al, 2019).

### 13. Wound healing assay, Matrigel invasion assay, Fluorescent gelatin degradation assay, Invadopodia formation assay

For determination of the cellular motility and invasive properties *in vitro* wound healing (O’Connell et al., 2008), matrigel invasion towards 20% FBS as chemoattractant followed by the crystal violet staining of invaded cells (Makowiecka et al., 2016), fluorescent gelatin degradation in the presence of 5% FBS after overnight starvation (Makowiecka et al., 2016) and invadopodia formation assays (Makowiecka et al., 2016) were prepared according to the established protocols. The wound gap was photographed using the Olympus ix51 inverted fluorescence microscope after 24 h and 48h from scratch. Wound width was measured by use of the QuickPHOTO MICRO 3.0 software. For the fluorescent gelatin degradation assay purpose, 80 ng/ml of rhWNT5A was used during 16 h of cells incubation on the coverslips coated with gelatin-Oregon Green conjugate (G13186, Thermo Fisher Scientific). Alexa Fluo 594 phalloidin (A12381, Thermo Fisher Scientific) and TO-PRO-3 Iodide (642/661) were employed for the cells visualization on confocal microscopy platform Leica TCS SP8. For invadopodia formation assay, immunofluorescence imaging protocol employing phalloidin and anti-cortactin antibody was performed. Invadopodia – as structures double positive for F-actin and cortactin staining, were quantified for tested cell lines and conditions and presented as number of invadopodia per one cell. Two independent repetitions were performed.

### 14. Colony formation assay

To assess an ability to colony formation in the presence of 0.3 μM vemurafenib, 300 of melanoma cells were plated onto 24-well plate and were subsequently cultured for seven days.

After that time, medium was removed and colonies were washed in PBS, fixed in the ice-cold methanol for 30 min and stained in the 0.5% crystal violet in 25% methanol. After washing and drying, bound crystal violated was eluted with 10% acetic acid and absorbance at 590 nm was measured on Tecan Sunrise plate reader. Result were normalized to the non-treated A375 wild type results.

### 15. Software, statistics

Statistical significance was confirmed by two-tailed paired or unpaired Student’s t tests. Statistical significance levels were defined as *P < 0.05; **P < 0.01; ***P < 0.001, ****p<0.0001. All statistical details including number of biological or technical replicates can be found in each figure legend. Statistical analysis and data visualization were performed in GraphPad Prism 8.0 software. Graphs are presented with error bars as ± SD if not stated differently in the figure legends.

## Acknowledgments

We would like to thank Lucie Nesvadbová, Lenka Bryjová, Pavlína Žofka Mrhálková, Lenka Doubková and Naďa Bílá for excellent assistance and also to Tomáš Bárta for technical support.

## Funding

The work was supported by the Czech Science Foundation grant GX19-28347X. CIISB, Instruct-CZ Centre of Instruct-ERIC EU consortium, funded by MEYS CR infrastructure project LM2018127, is gratefully acknowledged for the financial support of the measurements at the CEITEC Proteomics Core Facility. Computational resources for proteomics data processing were supplied by the project “e-Infrastruktura CZ” (e-INFRA LM2018140) provided within the program Projects of Large Research, Development and Innovations Infrastructures.

## Author contributions

TR, LK, ZZ, KS and VB designed the experiments and analyzed the data. TR, MN, KG, OVB, KAR, MP, RV, TG, KK, LD, LK, DP and KS performed the experiments. TR and VB wrote the manuscript. All authors discussed the results and commented on the manuscript.

## Competing interests

The authors declare that there is no conflict of interest regarding the publication of this article.

## Figures, supplements, data sources and tables list

Figure 1. RNF43 interactome is enriched with the Wnt Planar Cell Polarity pathway

Figure 1 Supplementary table 1 - BioID - RNF43 interactors

Figure 1 Supplementary table 2 - gProfiler GO terms analysis

Figure 2. RNF43 interacts with WntPCP components.

Figure 2 Source Data Figure 2 figure supplement 1.

Figure 2 figure supplement 1 Source Data

Figure 3. Mechanism of Wnt PCP inhibition by RNF43

Figure 3 Source Data

Figure 3 figure supplement 1.

Figure 3 figure supplement 1 Source Data

Figure 3 figure supplement 2.

Figure 4. RNF43 in melanoma.

Figure 4 Source data

Figure 4 figure supplement 1.

Figure 4 figure supplement 1 Source data

Figure 4 figure supplement 2.

Figure 4 figure supplement 2 Source Data

Figure 5. RNF43 inhibits WNT5A dependent invasive properties of human melanoma.

Figure 5 Source data

Figure 5 figure supplement 1.

Figure 5 figure supplement 2.

Figure 5 figure supplement 3.

Figure 5 figure supplement 4.

Figure 6. RNF43 overexpressing melanoma cells fail to develop resistance to bRAF inhibition.

Figure 6 Source Data

Table 1 – Cloning and mutagenesis primers

Table 2 – Antibodies

Table 3 – qPCR primers

Supplementary Table 1-Sequencing of the CRISPRCas9 derived cell lines

**Figure 2 figure supplement 1.**
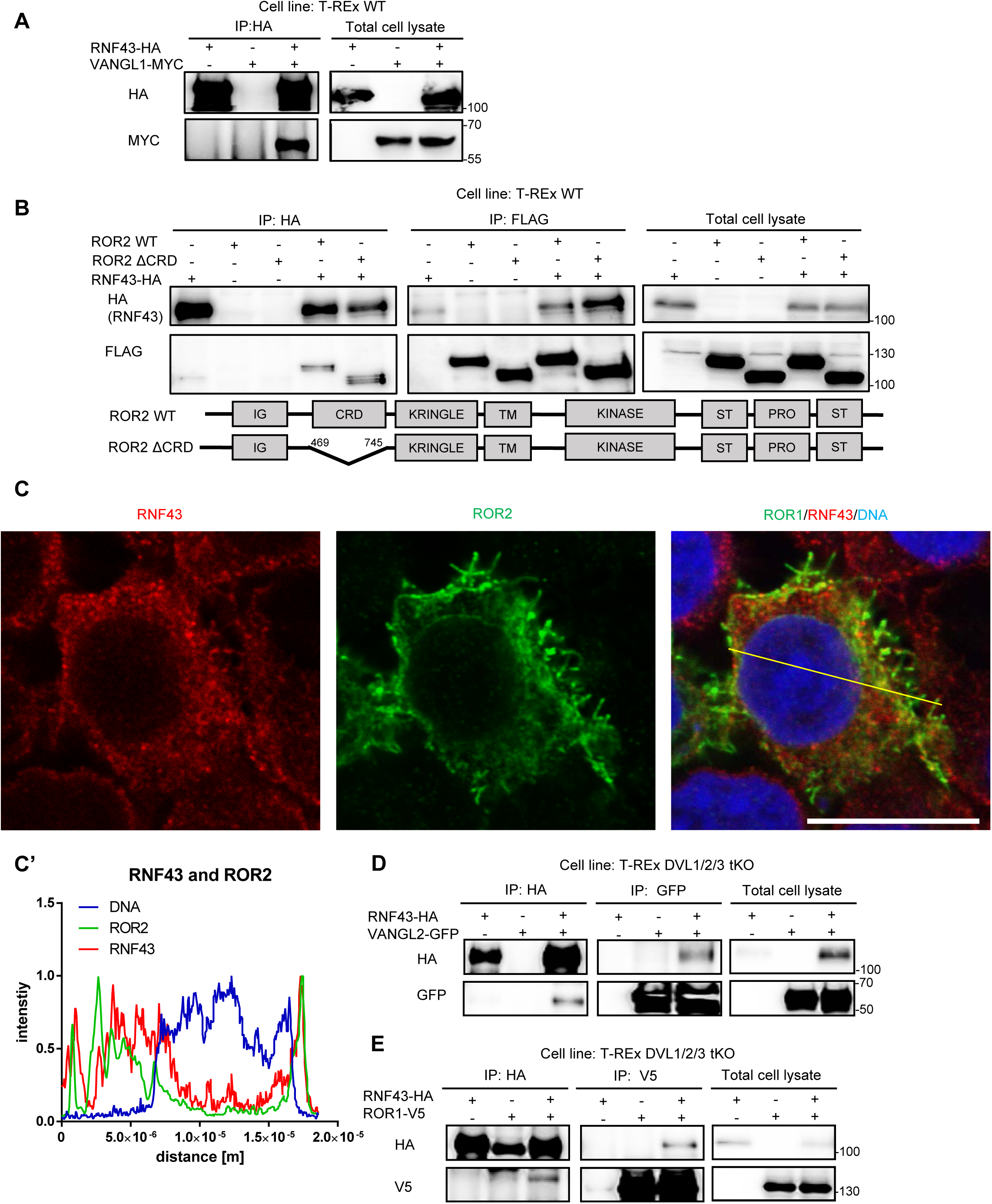
RNF43 interacts with Wnt/PCP components. **A**. RNF43 interacts with VANGL1. VANGL1-Myc was co-immunoprecipitated in the HA pull-down, prepared from lysate of Hek293 T-REx cells transiently overexpressing RNF43-HA and VANGL1-Myc, but not from the lysate containing only VANGL1-Myc overexpressed transgene, N=3. **B**. RNF43 interacts with the ROR2 in the CRD domain dispensable manner. Wild type ROR2 and ΔCRD-ROR2 mutant were detected in HA and FLAG pull downs, prepared from lysates of the Hek293 T-REx cells transiently overexpressing RNF43-HA and ROR2-FLAG or ΔCRD-ROR2, N=3. **C**., **C’**. Exogenous ROR2 (antiFLAG, green) colocalizes with the RNF43-BirA*-HA (anti-HA, red) in the TetON Hek293 T-Rex cells. DNA was visualized by TO-PRO-3 Iodide. Scale bar represents 25 μm. Co-localization of ROR2 and RNF43 was analyzed utilizing histograms (C’) of red, green and blue channels along selection (yellow line). Data is present in the Figure 2 figure supplement 1 Source Data. **D**. RNF43 interacts with the VANGL2 in the absence of all three Disheveled isoforms. RNF43 binding to VANGL2 in the DVL1/2/3 deficient cells was confirmed in the two-directional co-IP assay, N=3. E. Interaction between ROR1-V5 and RNF43-HA is preserved in the DVL1-3 null cells. ROR1 was detected in the HA pull-down and RNF43 in the V5 immunoprecipitation. T-REx DVL1/2/3 tKO cells were transfected with highlighted plasmids, N=3.

**Figure 3 figure supplement 1.**
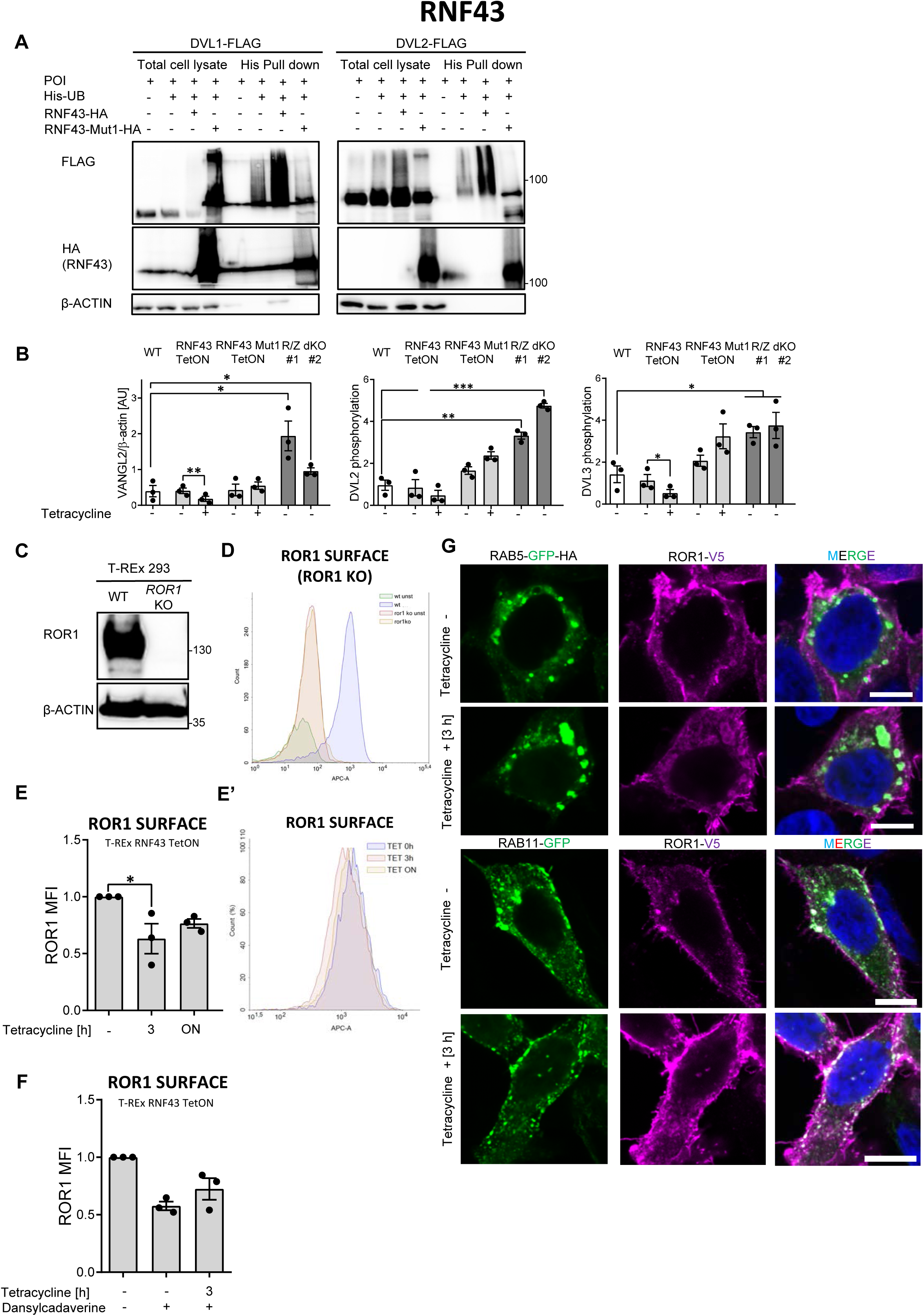
Mechanism of Wnt PCP inhibition by RNF43. **A**. DVL1 and DVL2 are ubiquitinylated by the E3 ubiquitin ligase RNF43, but not by its enzymatically inactive mutant (RNF43Mut1). Hek293 T-REx cells were transfected with plasmid encoding His-tagged ubiquitin, DVL1-FLAG or DVL2-FLAG and wild type or Mut1 RNF43 constructs and subjected to His-tag pull down and subsequent Western blotting. N=3. **B**. Quantification of Western blots from Fig. 3B. unpaired two-tailed t-test, *p<0.05; **p<0.01, ***p<0.001, N=3. **C**. Western blotting showing the lack of ROR1 protein in the T-REx 293 *ROR1* KO cell lines. **D**. T-REx *ROR1* KO line was used for validation of the ROR1-APC antibody used for the flow cytometric determination of the ROR1 cell surface level. **E**., **E’** Analysis of cell surface ROR1 in the T-REx RNF43 TetON cell line. ROR1 was internalized in the HA-positive (RNF43) cells population upon 3 hours tetracycline treatment, p= 0.0486, N=3. ROR1-APC flow cytometry histogram is shown (E’). **F**. Dansylcadaverine blocked RNF43-mediated effect on the ROR1 cell surface effect in the T-REx 293 RNF43 TetON cells, N=3. **G**. Immunofluorescence imaging of enhanced ROR1 (anti-V5, magenta) colocalization with marker of early endosomes RAB5 (GFP, green) after 3 hours of tetracycline treatment in the T-REx RNF43 TetON cell line. RAB11 (GFP, green) was recruited to the ROR1 at plasma membrane upon overnight tetracycline exposition. DNA was visualized by Hoechst 33342 (blue). Scale bars represent 10 µm. Cells were treated 24h post-transfection for indicated time points. Images representing RAB5 control and 3 h tetracycline treatment together with RAB11 control and tetracycline 3h conditions are presented in the Figure 3 figure supplement 2. Data presented in the B, E and F is presented in the Figure 3 figure supplement Source Data file.

**Figure 3 figure supplement 2.**
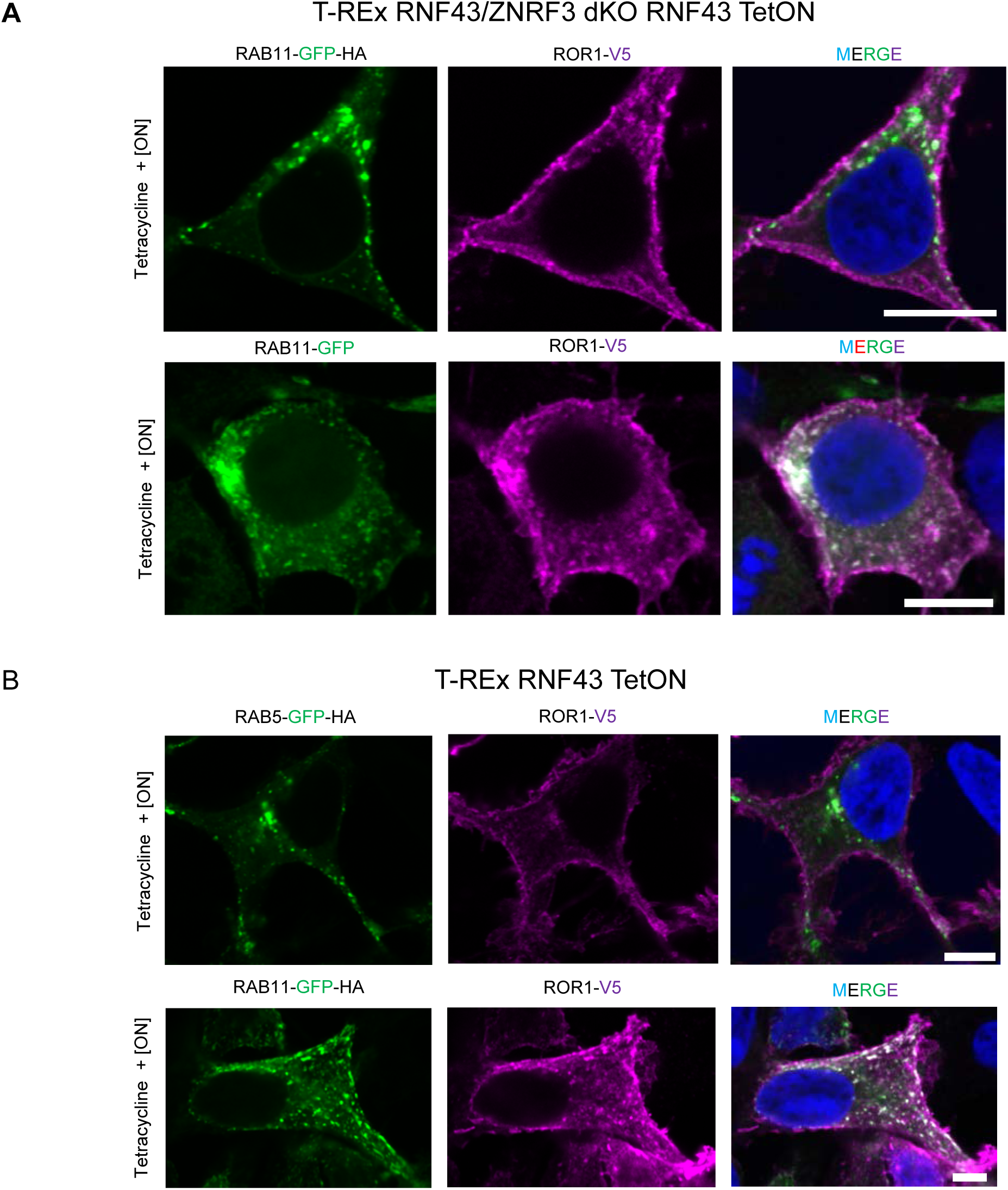
Mechanism of Wnt PCP inhibition by RNF43. **A**., **B**., Confocal imaging of the inducible T-REx RNF43/ZNRF3 dKO (A) and T-REx WT RNF43 TetON (B) and transfected with plasmids encoding ROR1-V5 (anti-V5, magenta) and RAB11-GFP-HA (GFP, geen). 24 hours post transfection, cells were treated with tetracycline for indicated time and then PFA fixed, permeabilized and stained for ROR1 (V5) and nuceli (DAPI, blue) detection. Scale bars represent 10 μm. Other tetracycline time points are presented in the Figure 3 and Figure 3 figure supplement 1.

**Figure 4 figure supplement 1.**
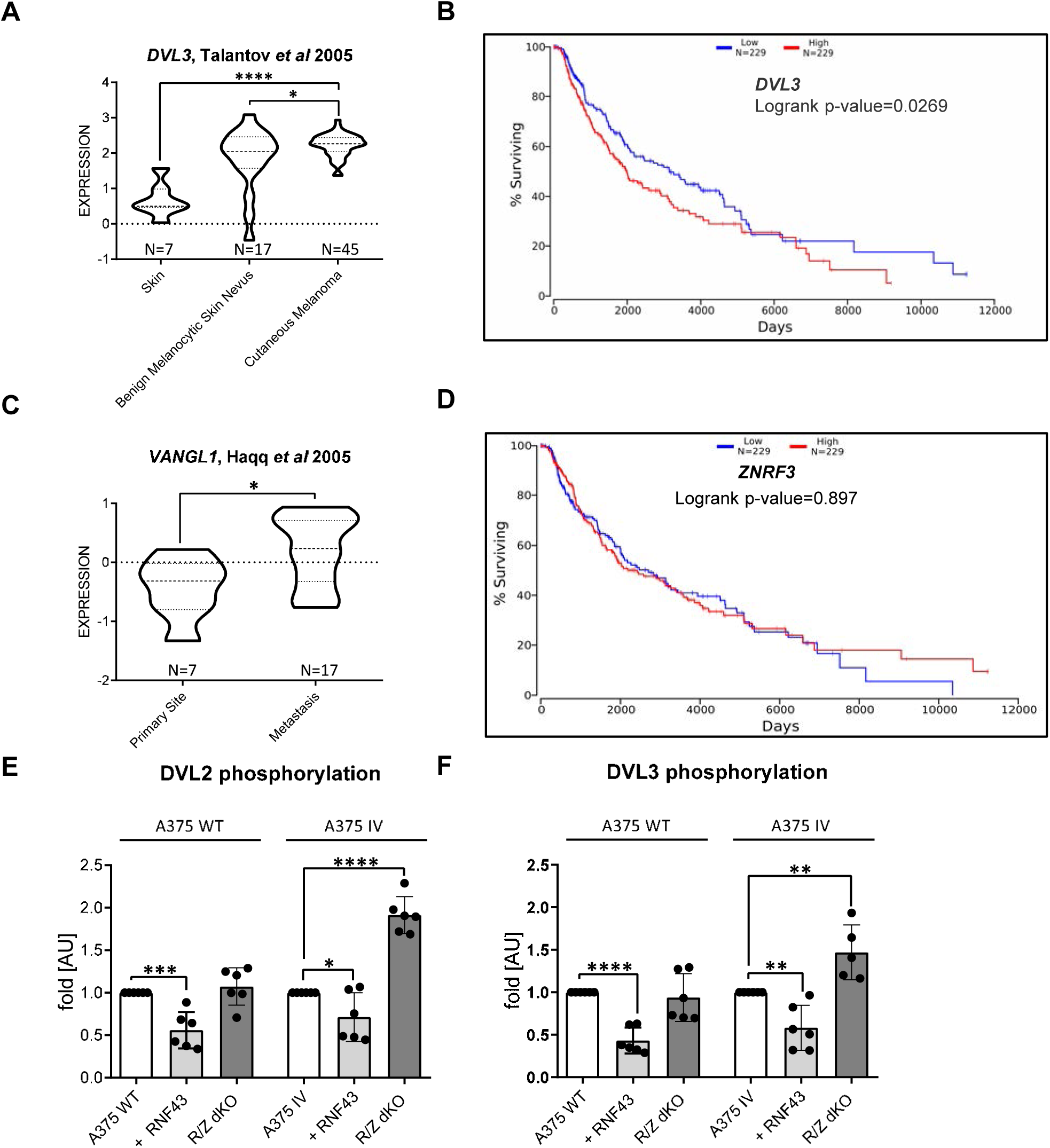
RNF43 in melanoma. **A**. *DVL3* expression level is elevated in human melanoma, unpaired two-tailed t-test, *p = 0,0159, ****p<0.0001. **B**. High expression of *DVL3* is a negative prognostic factor (50% lower and 50% upper percentiles). Logrank p-value=0.0269 **C**. *VANGL1* is more expressed in the metastasis than in primary melanoma, unpaired two-tailed t-test, p=0.0241. **D**. *ZNRF3* gene expression has no impact on melanoma patients survival. **E**., **F**. Quantification of Western blots presented in the Fig. 4G. Exogenous RNF43 expression blocked in both tested cell lines DVL2 (E) DVL3 (F) phosphorylation dependent shifts. CRISPR/Cas9 mediated knock-out of *RNF43*/*ZNRF3* resulted in the more activated DVL2 and DVL3 isoforms in case A375 IV cell line. Data were normalized to 1 for the parental cell lines values, unpaired two-tailed t-test: *p < 0.05, **p < 0.01, ****p < 0.0001, N=6 (F – A375 IV R/Z dKO N=5). Total DVL2 and DVL3 protein levels accompanied by *WNT5A, RNF43, ZNRF3, DVL2* and *DVL3* genes expression analysis is present in the Figure 4 figure supplement 2. Data used in the A, C, E and F is shown in the Figure 4 figure supplement 1 Source data.

**Figure 4 figure supplement 2.**
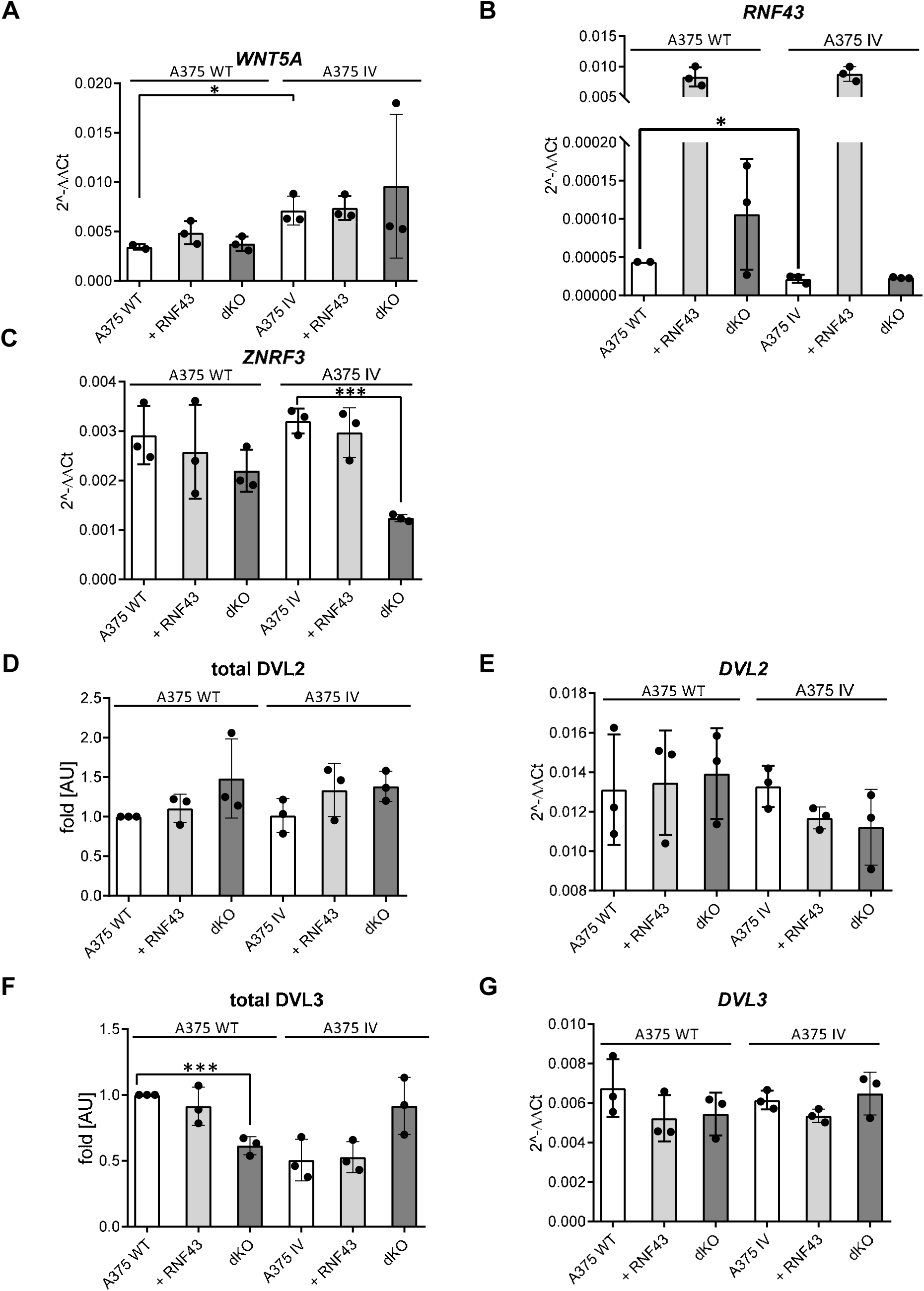
RNF43 in melanoma. **A**., **B**., **C**. RT-qPCR results – expression of the *WNT5A* (A), *RNF43* (B) and *ZNRF3* (C) genes was analyzed in the tested melanoma cells and presented as 2^−ΔΔCt^ ± SD, two tailed t-test: *p < 0.05, N=3 (A375 WT N=2 for A and B). Relative expression level was normalized to the *B2M* and *GAPDH* genes expression **D**., **E**. Western blot quantification results (Fig. 4G.) showing not affected by RNF43 overexpression or *RNF43*/*ZNRF3* knockout total level of DVL2 (D) and DVL3 (F). DVL3 protein level decreased only in the case of A375 WT R/Z dKO (D), unpaired two-tailed t-test: ***p < 0.001, N=3. Results were normalized to the A375 WT values. **E**., **G**. RT-qPCR analysis of *DVL2* (E) and *DVL3* (G) genes expression in A375 WT, A375 IV cell lines and their derivates. Relative expression level was normalized to the *B2M* and *GAPDH* genes expression values and presented as 2^−ΔΔCt^ ± SD. Data usied in the A – G is present in the Figure 4 figure supplement 2 Source Data.

**Figure 5 figure supplement 1.**
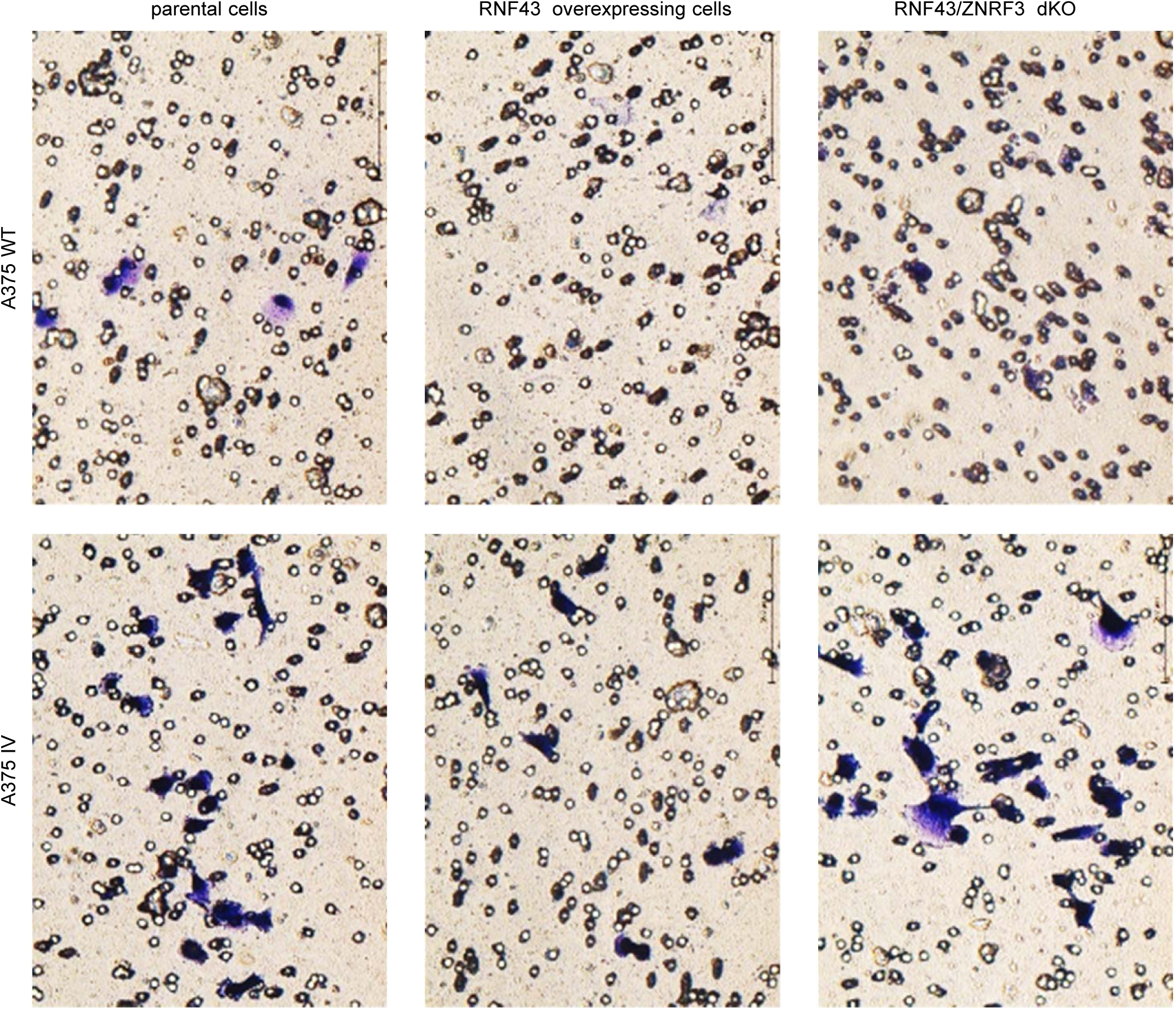
RNF43 inhibits Wnt5a dependent invasive properties of human melanoma. Representative photos of Matrigel invasion assay after crystal violet staining. Results are present in the Fig.5C.

**Figure 5 figure supplement 2.**
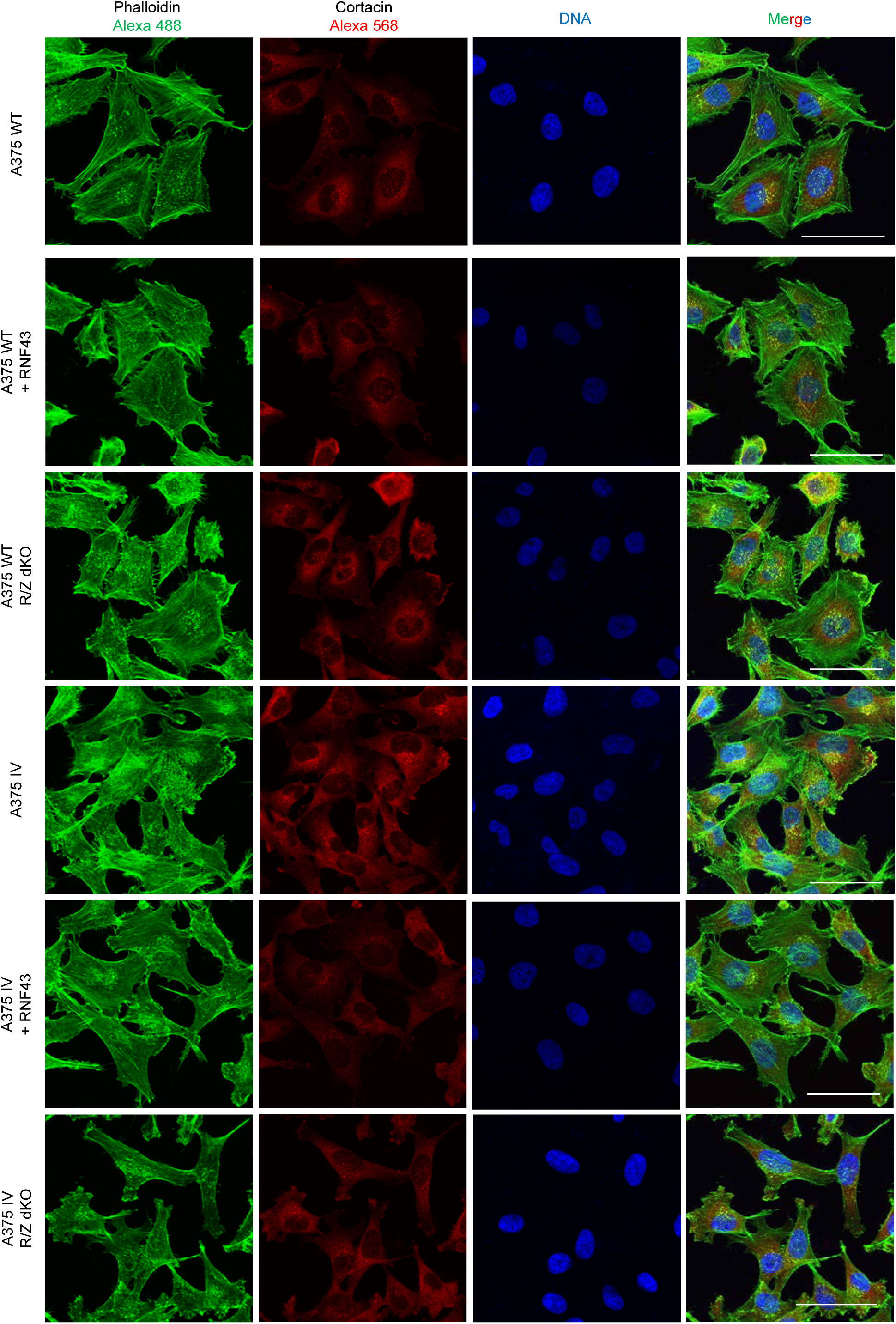
RNF43 inhibits Wnt5a dependent invasive properties of human melanoma. Confocal imaging of the A375 WT, A375 IV and RNF43 overexpressing and *RNF43*/*ZNRF3* double knock-out cell lines modifications. Cells were PFA fixed, Triton X-100 permeabilized and stained for cortactin by antibody (secondary antibody Alexa 568, red) F-actin using fluorescent phalloidin conjugate (Alexa 488, green) and TO-PRO-3 Iodide for DNA visualization (blue). Number of double positive puncta in single cells were quantified using imageJ software. Scale bars represent 100 μm. Results are present in the Fig.5D.

**Figure 5 figure supplement 3.**
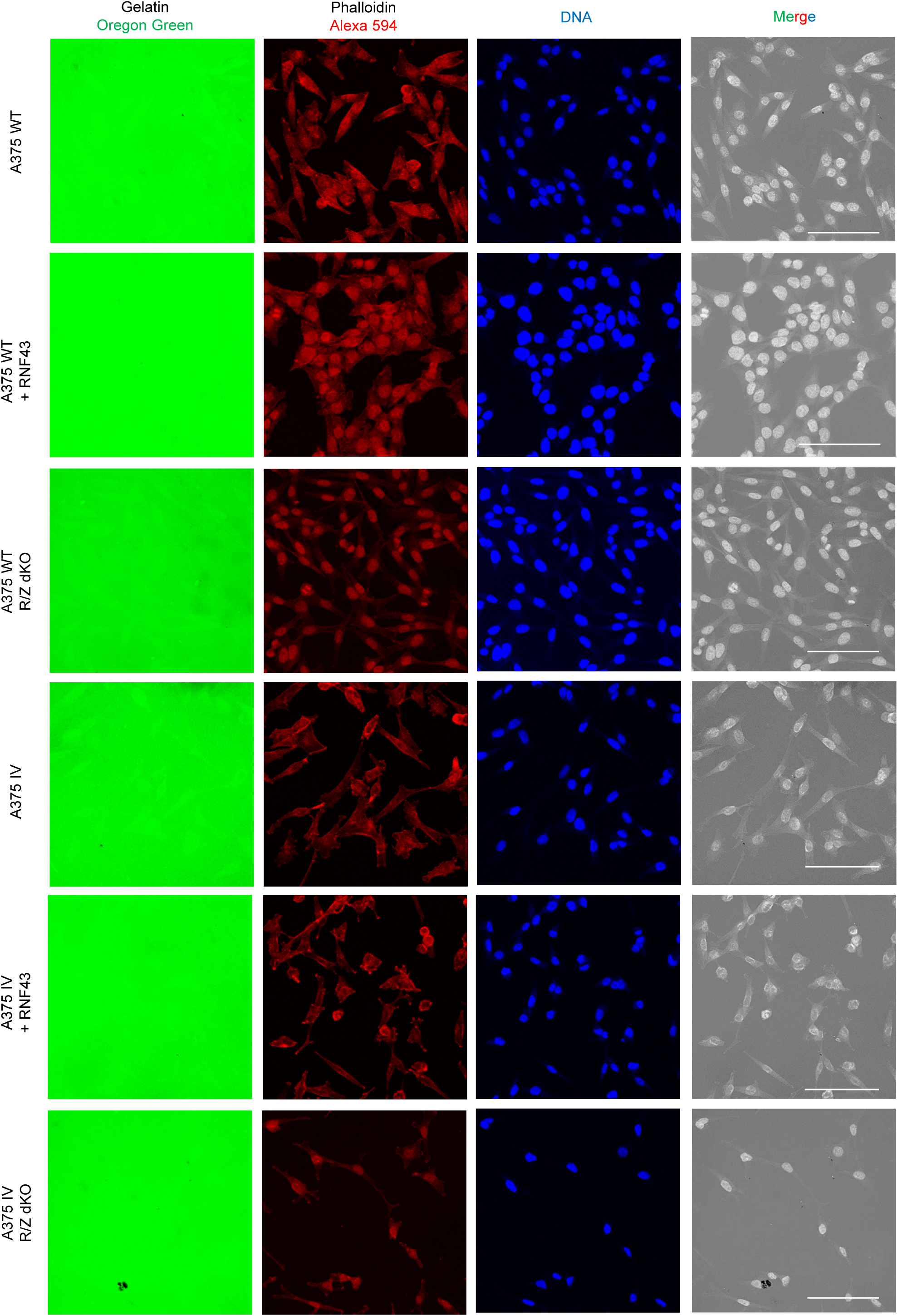
RNF43 inhibits Wnt5a dependent invasive properties of human melanoma. Confocal imaging of gelatin degradation assay without (Figure 5 figure supplement 3) and after (Figure 5 figure supplement 4) rhWNT5A treatment. Serum starved cells were plated onto gelatin-Oregon Green (green) coated coverslips and incubated for 24 hours.Fixed cells were stained with phalloidin-Alexa 594 for F-actin visualization (red) and TO-PRO-3 Iodide for nuclei (blue). Foci showing gelatin degradation are marked. Scale bars represent 50 μm. Experiment was repeated three times. Results are present in the Fig.5D.

**Figure 5 figure supplement 4.**
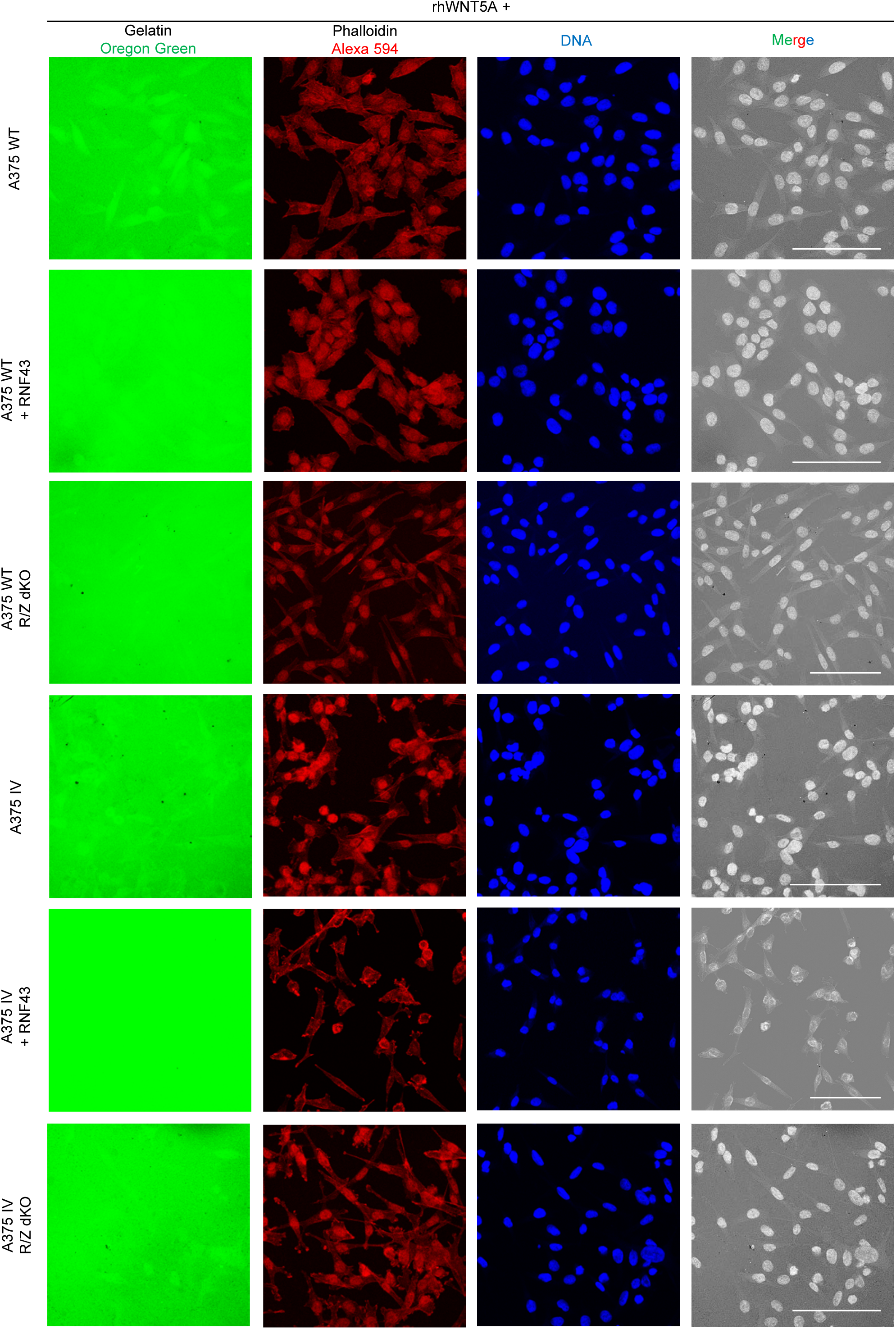
RNF43 inhibits Wnt5a dependent invasive properties of human melanoma. Confocal imaging of gelatin degradation assay without (Figure 5 figure supplement 3) and after (Figure 5 figure supplement 4) rhWNT5A treatment. Serum starved cells were plated onto gelatin-Oregon Green (green) coated coverslips and incubated for 24 hours.Fixed cells were stained with phalloidin-Alexa 594 for F-actin visualization (red) and TO-PRO-3 Iodide for nuclei (blue). Foci showing gelatin degradation are marked. Scale bars represent 50 μm. Experiment was repeated three times. Results are present in the Fig.5D.

